# Inference of fitness landscapes with heterogeneous patterns of epistasis across sites

**DOI:** 10.64898/2026.06.25.734428

**Authors:** Carlos Martí-Gómez, David M. McCandlish

**Affiliations:** Simons Center for Quantitative Biology, Cold Spring Harbor Laboratory, Cold Spring Harbor, NY, 11724

**Keywords:** fitness landscape, epistasis, Gaussian process

## Abstract

Fitness landscapes provide a framework for understanding how genetic variation shapes evolutionary outcomes. Although these landscapes were long treated as abstract conceptual objects, recent advances in genetic engineering and high-throughput phenotyping have enabled the empirical measurement of phenotypic values across large combinatorial sequence spaces. These developments create a need for statistical frameworks that can summarize, infer, and interpret fitness landscapes in the presence of complex genetic interactions. Here, we introduce a framework for summarizing the structure of genetic interactions across sites based on the average squared local *k*-way epistatic coefficients between mutations at different subsets of sites, and derive the precise manner in which the variance in these local *k*-way epistatic coefficients across backgrounds relates to epistasis of orders higher than *k*. These statistics can be computed exactly for complete combinatorial landscapes and are related to classical statistics in the fitness landscape literature. Moreover, they can be estimated from empirical correlations when data are incomplete or noisy, and used to define an empirical Bayes prior for fitness landscape inference that differentially penalizes interactions involving different subsets of sites. We apply this inference method to diverse high-throughput protein and RNA combinatorial mutagenesis datasets and find that fitness landscapes often show highly structured patterns of genetic interactions across positions. Finally, we use this model to infer a fitness landscape for a dynamic self-splicing intron comprising 65,536 genotypes, and describe in detail the main genetic interactions that shape the structure of this landscape and how they relate to the underlying molecular mechanism. Together, these results provide new tools for summarizing and modeling complex fitness landscapes, and for linking large-scale empirical data to the mathematical theory of fitness landscapes.

## Introduction

The fitness landscape is a fundamental concept in evolutionary biology and genetics. First introduced by Wright (1932), it describes the mapping between genotypes and their associated fitness. The structure of a fitness landscape, including the number, distribution, and connectivity of fitness peaks, plays a crucial role in how populations evolve and diverge over time (Kauffman and Levin 1987; Kondrashov et al. 2002; Gavrilets 2004; Weinreich *et al*. 2006; McCandlish 2011; De Visser and Krug 2014; Fragata et al. 2019; Manrubia et al. 2021; Bank 2022; Johnson et al. 2023). Understanding the mapping from genotype to fitness is not only important for explaining and predicting evolution, but also has critical applications in cancer and human disease (Moore and Williams 2009; Dasari et al. 2021) as well as plant and animal breeding (De Los Campos et al. 2013; Sackton and Hartl 2016; Soyk et al. 2020; Dwivedi et al. 2024). However, despite its importance, characterizing this mapping is inherently challenging due to the high dimensionality of sequence space. Because the number of possible sequences grows exponentially with sequence length, such high-dimensional fitness landscapes are often summarized by computing low-dimensional summary statistics that characterize the ruggedness or structure of the landscape, e.g. the number of local optima, lengths of adaptive walks, and the number of alternative local optima accessible from starting genotype (Kauffman and Levin 1987; Szendro *et al*. 2013; Ferretti *et al*. 2018). Another common approach is to characterize how mutational effects change across genetic backgrounds, for instance by quantifying the average magnitude of local epistatic coefficients (Zhou and McCandlish 2020) or by measuring the correlation of mutational effects between genotypes separated by increasing numbers of mutations (Weinberger 1990; Stadler 1996; Neidhart *et al*. 2013; Bank *et al*. 2016; Ferretti *et al*. 2016).

Historically, the scarcity of comprehensive experimental data has motivated the development of theoretical and computational models of fitness landscapes, which have provided a framework for understanding how summary statistics behave across different classes of landscapes. One approach is to consider families of fitness landscapes drawn from a probability distribution, generally known as random field models (Kauffman and Levin 1987; Stadler and Happel 1999). Classical examples include the House of Cards model (Kingman 1978), the NK model (Kauffman and Levin 1987), and the Rough Mount Fuji landscape (Aita and Husimi 1998). Within these frameworks, one can compute the expected values of summary statistics and analyze how they depend on parameters that control the smoothness or ruggedness of the landscape (Kauffman and Levin 1987; Schmiegelt and Krug 2014; Neidhart *et al*. 2014; Hwang *et al*. 2018; Reddy and Desai 2021).

More recent efforts have increasingly turned toward the experimental characterization of empirical fitness landscapes by measuring growth rates or other measures of biological functionality for many different combinations of mutations (De Visser and Krug 2014; Fragata et al. 2019). The size of the first empirically reconstructed fitness landscapes was limited by the difficulty of engineering large numbers of genotypes and measuring their fitness experimentally (Khan et al. 2011; Chou *et al*. 2011; Flynn et al. 2013; Szendro et al. 2013; Ogbunugafor *et al*. 2016; Weinreich et al. 2018; Gao et al. 2022; Aguirre et al. 2023; Zebell et al. 2025). However, recent advances in high-throughput assays (Kinney et al. 2010; Fowler and Fields 2014; Kinney and McCandlish 2019) have substantially expanded this scope, enabling the parallel measurement of thousands to millions of genotypes. These techniques have been used to characterize fitness landscapes across a range of biological systems, including regulatory sequences (Noderer et al. 2014; Rosenberg *et al*. 2015; Bonde et al. 2016; Evfratov et al. 2017; Rabani et al. 2017; Wong et al. 2018; Baeza-Centurion et al. 2019; Kuo et al. 2020; Komarova et al. 2020; de Boer et al. 2020; Vaishnav et al. 2022; Liao et al. 2023; Westmann et al. 2024b,a; Kuo et al. 2025; Chattopadhyay et al. 2025; Agarwal et al. 2025), RNAs (Domingo *et al*. 2018; Bendixsen et al. 2019; Soo et al. 2021; Rotrattanadumrong and Yokobayashi 2022), proteins (O’Maille et al. 2008; Bank *et al*. 2016; Wu et al. 2016; Starr et al. 2017; Poelwijk et al. 2019; Lite et al. 2020; Jalal et al. 2020; Bryant et al. 2021; Somermeyer *et al*. 2022; Moulana et al. 2023; Papkou et al. 2023; Sundar et al. 2024; Zarin and Lehner 2024; Johnston et al. 2024; Faure et al. 2024b; Escobedo et al. 2025; Herrera-Álvarez et al. 2025), and genome-wide gene interactions (Bakerlee et al. 2022; Nguyen Ba et al. 2022; Matsui et al. 2022; N’Guessan et al. 2025). Despite these advances, contemporary datasets are often noisy and typically do not cover the full sequence space, so that a key challenge is to develop flexible statistical methods for inferring full fitness landscapes from empirical data without distorting the rich fitness landscape geometry revealed by these high-throughput measurements.

One powerful approach to address this problem is to combine theoretical models of fitness landscapes with empirical measurements by recasting these theoretical models as Bayesian priors for reconstructing complete landscapes from incomplete and noisy data (Zhou and McCandlish 2020; Chen et al. 2021; Zhou et al. 2022, 2025; Petti et al. 2025; Martí-Gómez et al. 2026b). Such an approach can leverage our mathematical understanding of these models to define prior distributions that confer the overall inference procedure with desirable properties. For example, Minimum Epistasis Interpolation defines a prior that depends on the average squared epistatic coefficients between all pairs of mutations (Zhou and McCandlish 2020), favoring reconstructions that are locally approximately additive, which results in reconstructions that can capture genetic interactions of all orders where data is abundant but extrapolates additively far from the data. As another example, Empirical Variance Component regression (Zhou et al. 2022) constructs a prior parametrized by the variance explained by genetic interactions of each possible order, resulting in reconstructions that accurately reflect how quickly the predictability of mutational effects decay in increasingly distant genetic backgrounds.

While Minimum Epistasis Interpolation and Empirical Variance Component regression can incorporate epistatic interactions of all orders, the corresponding priors are still only weakly informative in the sense that they are “isotropic” (Stadler 1996, 2002), i.e. the prior treats all sites and all mutations equally. However, in reality some sites and alleles are more influential and more likely to be involved in epistatic interactions than others (Weinreich *et al*. 2005; Kvitek and Sherlock 2011; Ferretti *et al*. 2016; Bank *et al*. 2016; Pokusaeva *et al*. 2019; Reddy and Desai 2021). Reddy and Desai (2021) recently proposed a new family of theoretical random field models whose parameters control the site-specific probability that mutations participate in genetic interactions, and these models were then extended by Zhou *et al*. (2025) to include allele-specific and mutation-specific parameters. By treating these site-, allele-, or mutation-specific parameters as hyperparameters of an informative Bayesian prior, the resulting model learns which mutations most strongly influence the predictability of other mutations, and incorporates this information when inferring the fitness landscape from data, achieving state-of-the-art predictive performance (Zhou *et al*. 2025). Nonetheless, these models still implicitly assume that the the propensity for a set of sites to interact is determined solely by these site-specific parameters, while empirical observations from both pairwise interaction models (Marks *et al*. 2012; Haldane *et al*. 2016, 2018) and the posterior distributions of models containing interactions of all orders (Chen *et al*. 2021; Martí-Gómez *et al*. 2026b) suggest that patterns of epistatic interaction are often sparse (Poelwijk *et al*. 2019) or modular (Rojas Echenique *et al*. 2019; Hwang *et al*. 2018).

Here, we present a method for fitness landscape inference incorporating a prior that can encode this type of highly structured tendency for specific sets of sites to interact with each other. We begin by proposing a simple approach to summarize the structure of genetic interactions of all orders by computing, for each pair of sites, the average squared local epistatic coefficient between mutation at those sites, which enables the identification of sets of sites involved in epistatic interactions of arbitrary order in complete combinatorial fitness landscapes. Based on these summary statistics, we define a new family of prior distributions that differentially penalize interactions involving different pairs of sites. The parameters of these priors can be estimated from empirical correlations in fitness values between sequences that differ at specific subsets of sites, and we then use the resulting priors to infer complete fitness landscapes from several empirical datasets. Finally, we apply our method to the fitness landscape of a dynamic self-splicing intron (Soo *et al*. 2021), and show how the higher-order interactions in this system have an interpretable structure wherein many aspects of the genetic architecture are systematically rewired depending on the nucleotide identities at one specific pair of sites.

## Results

### Epistatic coefficients for subsets of sites

In this section, our aim is to quantify the overall amount of epistasis between a specific pair of sites in an arbitrary fitness landscape *f*. We start by considering a simple two-locus biallelic fitness landscape and assuming that we know the fitness values of the four possible genotypes *f* _*AB*_, *f* _*Ab*_, *f*_*aB*_ and *f*_*ab*_. We can measure how much the effect of a mutation *A* → *a* in one locus changes in the presence of an additional mutation *B* → *b* at the other locus via the traditional double-mutant epistatic coefficient:3

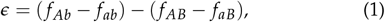

and we can use the squared epistatic coefficient *ϵ*^2^ to quantify the magnitude of epistasis in a manner that is independent of which physical allele is encoded as e.g.”A” or “a”. As we add a third locus, the epistatic coefficient between mutations *A* → *a* and *B* → *b* can be calculated in the presence of alleles *C* and *c* at this new locus, where these two different genetic backgrounds can be viewed as defining parallel faces in sequence space (Figure 1). Thus, one way of quantifying the extent of epistasis between sites 1 and 2 is by taking the average of the squared local epistatic coefficients defined on each of these two parallel faces. Similarly, we can quantify epistasis between sites 1 and 3 by averaging the local squared epistatic coefficients found on the corresponding pair of parallel faces, and we can quantify epistasis between sites 2 and 3 by averaging the squared local epistatic coefficients from the final remaining pair of faces (Figure 1).

**Figure 1.**
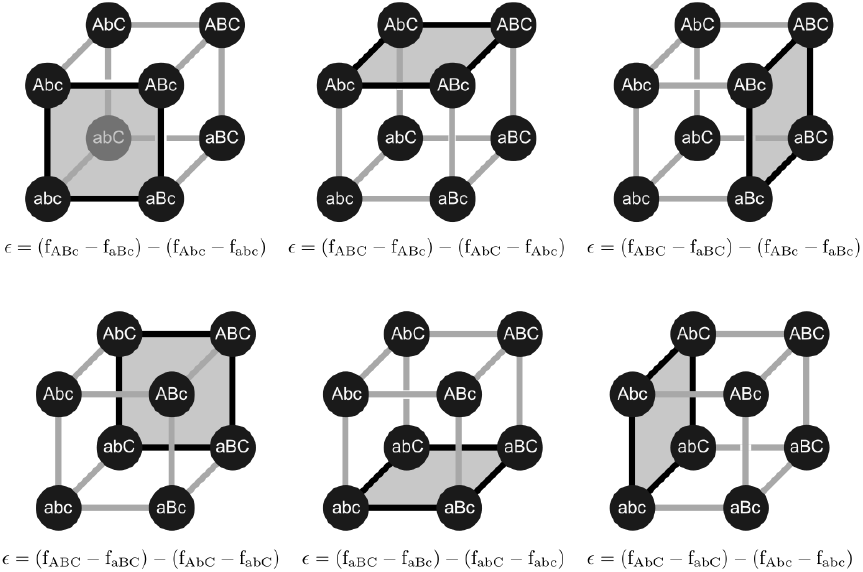
Local epistatic coefficients for mutations at different pairs of sites in a three-site bi-allelic fitness landscape. Local epistatic coefficients can be grouped into sets corresponding to the interaction of the same pair of mutations on different genetic backgrounds. Geometrically this corresponds to grouping epistatic coefficients on parallel sets of “faces”; here we show each set of parallel faces in its own column, with those corresponding to interactions between sites 1 and 2 in the first column, those corresponding to interactions between sites 2 and 3 in the middle column, and those corresponding to interactions between sites 1 and 3 in the last column.

We can generalize these statistics for larger fitness landscapes composed of *ℓ* sites with *α* alleles per site represented by a vector *f* of length *α*^*ℓ*^. In particular, epistasis between any pair of sites *i, j* can be quantified by computing the average squared local epistatic coefficient 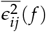, where the average is taken over every possible pair of mutations at those sites and across every possible genetic background in which they can be introduced. It is easy to see that the average of these quantities 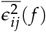 across all possible pairs of sites corresponds to the previously proposed average squared epistatic coefficient 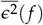 describing the overall local smoothness of a fitness landscape (Zhou and McCandlish 2020) (see Supplement). However, separately averaging these squared epistatic coefficients for each pair of sites provides a more granular description of epistasis, showing not only how much epistasis there is, but whether it is equally distributed across different pairs of sites or concentrated within specific subsets of sites.

For the special case of bi-allelic fitness landscapes, Ferretti *et al*. (2016) previously discussed the 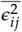 as part of their proposal of the statistic *γ*_*i j*_, which measures the correlation in the effects of mutations at site *j* before and after mutating site *i*. Here, we generalize the 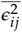 statistics to multi-allelic landscapes and to higher-order epistatic coefficients between any subset of sites *U* (see Supplement). Moreover, we show the relationship between these statistics and the total variance explained by interactions involving the sites in *U* of order equal or higher than |*U*|. Specifically, let Var^(*U*)^[*f*] be the total variance explained by |*U*|-way interactions among the sites in *U* as well as interactions of order greater than |*U*| that also involve all the sites in *U* (Reddy and Desai 2021; Martí-Gómez et al. 2026b). In the supplement, we show that this variance Var^(*U*)^[*f*] is proportional to the average squared local |*U*|-way epistatic coefficient 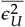:

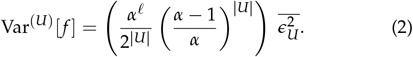

In addition, we show that the portion of this variance explained by genetic interactions of order strictly higher than |*U*| is proportional to the variance across backgrounds in |*U*|-way local epistatic coefficients between mutations at sites *U*:

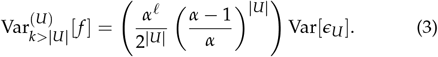

These results provide a quantitative link between the size and variance of local |*U*|-way epistatic coefficients involving a set of sites *U* and the amount of |*U*|-way and higher epistatic variance explained by that set of sites.

### Local epistasis regression

In the previous section, we defined a set of descriptive statistics 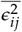 quantifying the magnitude of the local double-mutant epistatic coefficients when introducing mutations at a specific pair of sites *i* and *j*. While these statistics can only be evaluated for complete fitness landscapes, we can easily use them to construct a prior distribution for use in inferring complete fitness landscapes from noisy and incomplete data. We call the resulting Bayesian regression method Local Epistasis Regression.

In particular, we consider the prior

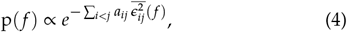

where the 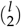 hyperparameters *a*_*ij*_ *>* 0 penalize the size of the local epistatic coefficients between each specific pair of sites *i* and *j*. This prior turns out to be an improper Gaussian prior that penalizes the size of local genetic interactions as specified by the *a*_*ij*_ but which places a flat prior over the non-epistatic component of the landscape.

In order to better understand the properties of this prior, and in particular why this prior allows epistatic interactions of all orders, it is helpful to re-express this new prior within a more general class of priors previously described by Zhou *et al*. (2022) and Martí-Gómez *et al*. (2026b). This more general prior is a Gaussian random field model parametrized by the variance explained by epistatic interactions between each possible subset of sites *U*. The prior as a whole is given by

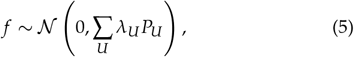

where *P*_*U*_ is the projection matrix into the subspace of landscapes composed solely of |*U*|-way interactions between the sites *U* 4 Local epistasis regression and we constrain the 2^*ℓ*^ hyperparameters *λ*_*U*_ to be non-negative, *λ*_*U*_ ≥ 0. This is the most general model in which the covariance between two sequences *x, x*′ only depends on the set of sites at which they differ, and the expected variance explained by |*U*|-way genetic interactions between a set of sites *U* is given by (*α* − 1)^|*U*|^ *λ*_*U*_ (see Supplement).

Turning back to our prior in terms of the *a*_*ij*_ and 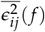 given in Equation 4, in the Supplement we show that this prior is equivalent to choosing

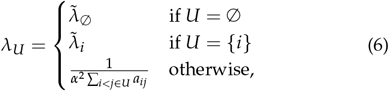

in the limit where 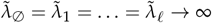. Since we assume *a*_*ij*_ *>* 0, we see that *λ*_*U*_ *>* 0 for |*U*| ≥ 2 indicating that our prior has positive variance for every *U* with |*U*| ≥ 2 and can thus fit genetic interactions of all orders. In practice, instead of taking the limit and working with an improper prior, we retain 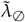 and 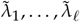. as additional hyper-parameters that provide control over the extent to which the non-epistatic component of the model is regularized. Finally, although *a*_*ij*_ penalizes the size of 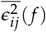, the expected value of 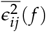 under the prior depends not only on *a*_*ij*_, but also on the *a*_*ik*_ and *a*_*jk*_ for all the *ℓ* − 2 other sites *k* (see Supplement).

In order to perform inference under this prior, we further assume that the experimental measurements *y* for a series of sequences *X* have normally distributed errors with known experimental variance *y*_*var*_ so that we can use standard Gaussian process results (Rasmussen and Williams 2008) to compute the closed form Gaussian posterior distribution over the complete fitness landscape *f* with mean

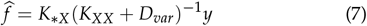

and covariance matrix

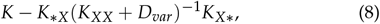

where *K*_*XX*_, *K* _* *X*_, *K*_* *X*_ are submatrices of *K* = ∑_*U*_ *λ*_*U*_ *P*_*U*_ indexed by sequences *X* and *, where * represents all possible sequences, and *D*_*var*_ is a diagonal matrix with the known experimental variances *y*_*var*_ along the diagonal. Naively evaluating and computing with these matrices becomes quickly impractical beyond a few thousand measurements. However, we can leverage the mathematical properties of the covariance matrix to derive efficient routines for computing matrix-vector products without explicitly constructing the matrices, which can be implemented as linear operators and used for solving large linear systems using iterative methods in *gpmap-tools* (see Supplement) (Martí-Gómez et al. 2026b).

The final remaining issue is how to choose the values for the 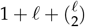 hyperparameters (specifically, 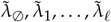 and the 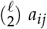). In principle, these hyperparameters can be specified based on prior knowledge, for example if certain sites are known to interact more strongly with each other such as contacting residues within a known structure. Here, however, we adopt an empirical Bayes approach and assume that the statistical properties of the measured sequences generalize to the full sequence space, allowing us to estimate these parameters directly from the data without needing any a priori knowledge about the structure of genetic interactions between sites. Practically speaking, because the covariance in fitness between two sequences under the prior depends only on the set of sites *D* at which they differ, we can compute the empirical correlation in measured fitness between all pairs of sequences differing exactly at a set of sites *D*, for each possible subset *D*. We then choose the hyperparameters so that the correlations implied by the prior match these empirical correlations as closely as possible, using a procedure known as kernel alignment (Wang *et al*. 2015; Zhou *et al*. 2022). This structure reduces the kernel alignment problem to a 2^*ℓ*^-dimensional weighted least squares problem, which can be solved efficiently under the constraint that the *a*_*ij*_, the 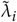 and 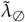 are all positive (see Supplement).

### Validation on simulated data

We first evaluate the performance of Local Epistasis Regression using simulated data to illustrate the advantages of this model in comparison with previously proposed approaches. Specifically, we define a random field model inspired by the expected interactions between 8 positions forming an RNA helix, where sites interact more strongly with their neighboring positions as well as with the sites with which they form base-pairs (Figure 2A). Figure 2B shows the correlation under this prior between pairs of sequences differing at every possible subset of sites *D* within each Hamming distance class and where lines represent distance classes that differ from each other at a single position e.g. *D*_1_ = {1, 2} and *D*_2_ = {1, 2, 3}. We see that while the correlation under the prior generally decays with increasing Hamming distance *d* = |*D*|, the correlation is not a strict function of *d* and instead varies based on the specific positions at which two sequences differ.

**Figure 2.**
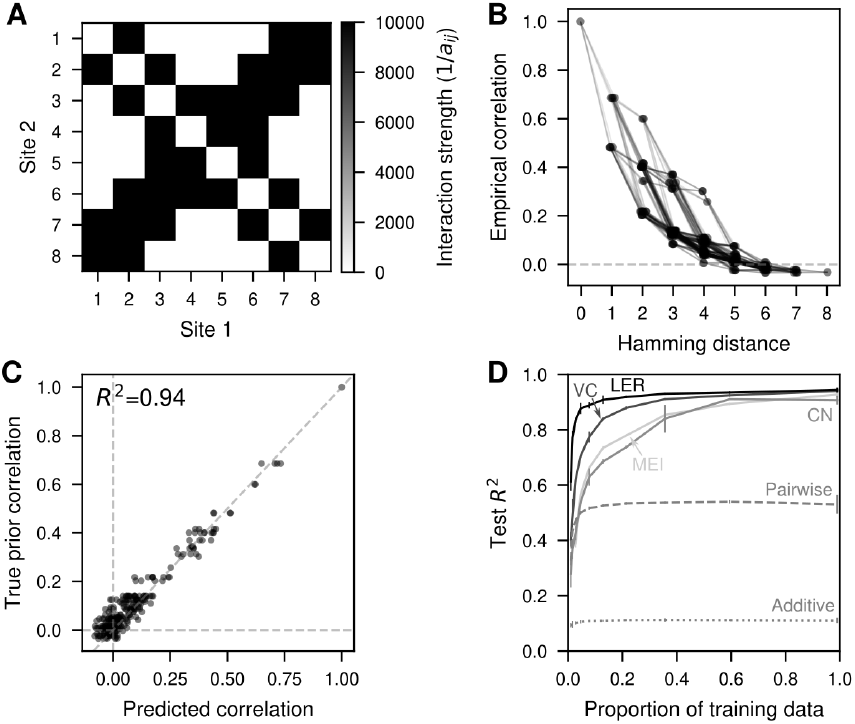
Validation of Local Epistasis Regression. (A) Heatmap representing the model hyperparameters as 1/*a*_*ij*_ for every pair of sites *i, j* highlighting the patterns of genetic interactions across sites under the prior. (B) Correlation under the prior for pairs of sequences that differ at each possible subset of sites *D* arranged according to the Hamming distance *d* = |*D*|. Each dot represents a single distance class *D* and are joined by lines whenever the distance classes differ by a single position from each other. (C) Comparison of the true correlation under the prior and the estimated ones using kernel alignment on a simulated dataset comprising 15% of the sequences from a fitness landscape drawn from the prior. (D) Predictive performance evaluated by the *R*^2^ between the predicted and the true fitness values of held-out test sequences when using different amounts of training data for different models (MEI: Minimum Epistasis Interpolation, VC: Variance Component regression, CN: Connectedness Model regression, LER: Local Epistasis Regression). Predicted values are the maximum a posteriori estimate given by each method, which is equal to the posterior mean 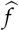. Error bars represent the standard deviation across 3 different random samples for each fraction of training data.

Having specified a family of random fitness landscapes, we then drew a specific fitness landscape from this prior distribution and evaluated the performance of the hyperparameter estimation procedure in recovering the ground truth hyperparameters in the presence of limited amount of data. In particular, we kept fitness values for only 15% of the sequences, computed the empirical fitness correlation between pairs of sequences for each class of differences *D* and estimated the hyperparameters for Local Epistasis Regression using kernel alignment. Figure 2C shows that we can accurately infer a prior that approximates the true correlation structure of the landscape using only a limited amount of data (*R*^2^ = 0.94).

Since our hyperparameter estimation procedure appeared to be working, we then evaluated the performance of the full Local Epistasis Regression procedure for different random samples of training data ranging from 1% to 99% of the full landscape and compare it with the predictive power of previously proposed models, including classical models like additive and pairwise interaction models, and models that allow interactions of all possible orders, including Minimum Epistasis Interpolation (Zhou and McCandlish 2020), Empirical Variance Component regression (Zhou *et al*. 2022) and a new implementation of Connectedness regression (Zhou *et al*. 2025) in *gpmap-tools* (Martí-Gómez *et al*. 2026b) that uses kernel alignment for hyperparameter inference (Figure 2D, see Supplement). Our results show that all models that allow higher order interactions can accurately reconstruct the true landscape given sufficient amount of data, unlike additive and pairwise interaction models. Moreover, by encoding information about which sites interact with each other and at which sites mutations combine more additively, Local Epistasis Regression outperformed the other methods for any amount of training data, and exhibited a particularly strong advantage for low amounts of training data.

### Learning the structure of genetic interactions from empirical data

In the previous section, we have shown how Local Epistasis Regression can be used to learn the structure of genetic interactions from incomplete and noisy data, and then used that information to make more accurate inference of complete combinatorial fitness landscapes using simulated data. Here, we investigate the performance of Local Epistasis Regression using data from diverse empirical fitness landscapes.

First, we used data from a high-throughput experiment evaluating the functionality of nearly every possible 5′ splice site sequence of the exon 7 in the Smn1 gene context (Wong *et al*. 2018; Zhou et al. 2022) (positions −3 through −1 at the exonic region and +2 through +6 at the intronic region are variable, with position +1 fixed as G, which is necessary for lariat formation during the splicing reaction). We start by computing the correlation between pairs of sequences differing at each possible combination of sites (Figure 3A). This analysis shows that the correlation between pairs of sequences depends, not only on the Hamming distance, but also on the specific combination of sites that are different. For example, pairs of sequences differing only at position +6 show a correlation of 0.83, whereas pairs of sequences that differ only at position +2 show a correlation of 0.12. Next, we estimated the hyperparameters of the prior for our Local Epistasis Regression model and show that the estimated prior can recapitulate the observed correlations almost perfectly (Figure 3B). Figure 3C displays the estimated strength of the penalization for local epistatic coefficients for mutations between each possible pair of sites, showing a rich structure of interactions between positions. First, we find that mutations interact more strongly with mutations at neighboring positions, as expected from its recognition mechanism by the U1 snRNA via base-pair complementarity (Wong *et al*. 2018) because the energetic contributions of a basepair to the thermodynamic stability of an RNA helix depend strongly on the adjacent basepairs (Borer *et al*. 1974). There are some exceptions to this general interaction pattern between positions: i) mutations at positions +2 and +5 tend to interact more strongly with each other than expected under the neighbor interaction model, as previously shown in the 5′splice site fitness landscape inferred from their frequencies in the human genome (Chen *et al*. 2021), and ii) the effects of mutations at position +6 are expected to combine more additively with the effects of other mutations. Interestingly, while mutations at position +2 show strong interactions with mutations at − 1 and +5, mutations at − 1 and +5 are expected to combine nearly additively, a pattern of interaction that cannot be captured by simpler random field models such as the Connectedness Model (Reddy and Desai 2021; Zhou *et al*. 2025). Finally, we evaluated the performance of Local Epistasis Regression when predicting the fitness of held out sequences for different amounts of training data (Figure 3D) and found that, given sufficient training data, predictions are more accurate than under all previously proposed models.

**Figure 3.**
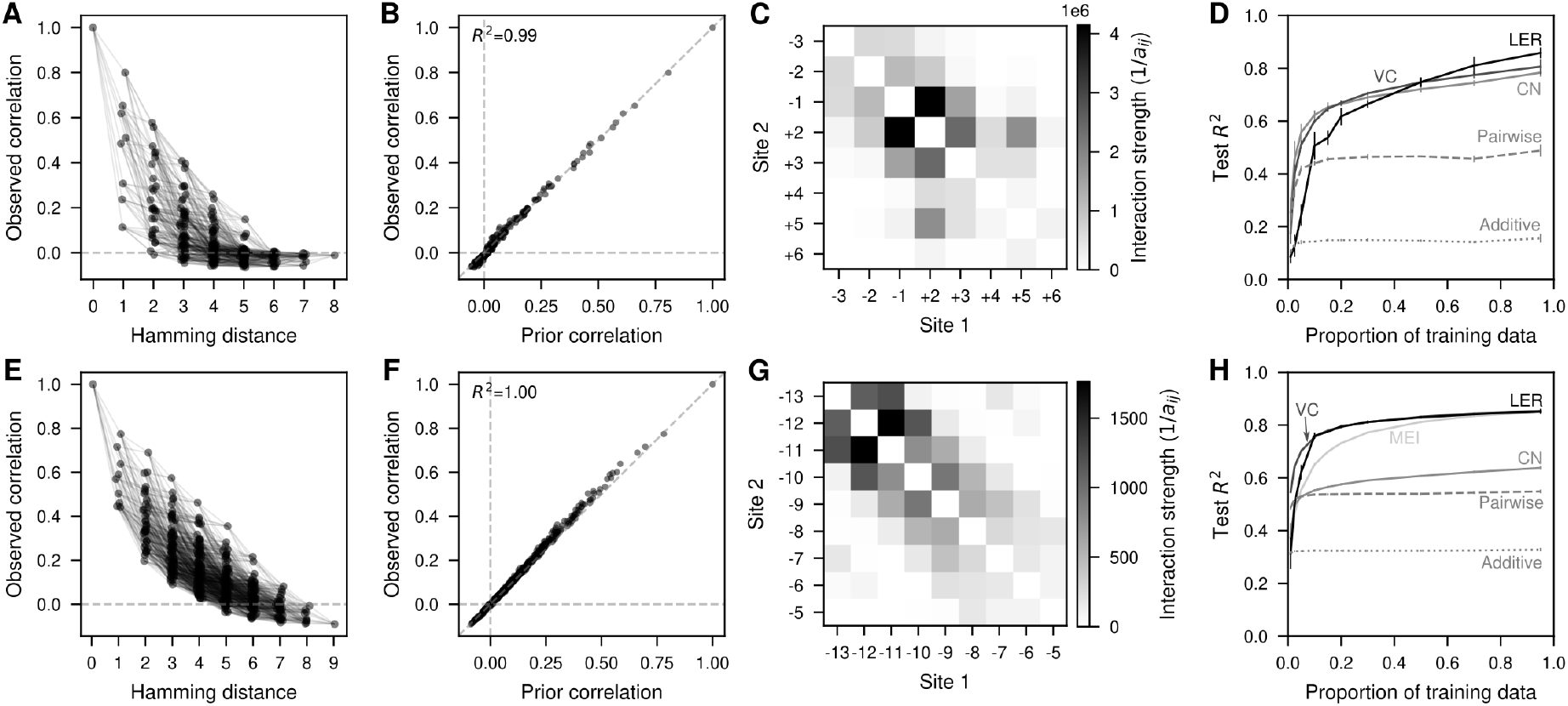
Application of Local Epistasis Regression to estimate the structure of epistatic interactions across sites in empirical fitness landscapes. (A,E) Correlation in the measured fitness values for pairs of sequences differing at each possible subset of sites *D* arranged according to the Hamming distance *d* = |*D*|. Each dot represents a single distance class *D* and are joined by lines whenever the distance classes differ by a single position from each other. (B,F) Comparison of the observed correlation values in the data and the values under the estimated prior ones using Local Epistasis Regression for every possible distance class *D* (each dot represents a different *D*). Correlations were estimated using 80% of the data for training. (C,G) Heatmap representing the inferred model hyperparameters as 1/*a*_*ij*_ for every pair of sites *i, j* highlighting the patterns of genetic interactions across sites under the prior. (D,H) Predictive performance evaluated by the *R*^2^ between the predicted and the measured fitness of held-out test sequences when using different amounts of training data for different models (MEI: Minimum Epistasis Interpolation, VC: Variance Component regression, CN: Connectedness Model regression, LER: Local Epistasis Regression). Predicted values are the maximum a posteriori estimate given by each method, which is equal to the posterior mean 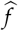. Error bars represent the standard deviation across 3 different random samples for each fraction of training data. Each row represents a fitness landscape: Smn1 exon 7 5′ splice site (A,B,C,D); dmsC Shine-Dalgarno sequence (E,F,G,H).

Second, we used data from a high-throughput experiment measuring the translational activity of over 250,000 9-nucleotide sequences at the Shine-Dalgarno sequence in the 5′UTR of the dmsC gene in *E. coli* (Kuo *et al*. 2020). As before, the correlation in the measurements between pairs of sequence depends not only on the Hamming distance between them, but also on the specific combination of sites at which they differ (Figure 3E). These empirical correlations can be accurately captured by the Local Epistasis Regression prior under the estimated hyperparameters (Figure 3F). The inferred hyperpameters recapitulate the previously characterized pattern of genetic interactions between mutations at different pairs of sites for this landscape, where sites interact more strongly with other sites within a 4-nucleotide window (Figure 3G) (Martí-Gómez *et al*. 2026b). While the Shine-Dalgarno sequence is also recognized via basepair complementarity, in this case with the 16S rRNA (Shine and Dalgarno 1975), the observed pattern of interaction between positions can be explained by the ability of the 16S rRNA to bind the target sequence at different registers relative to the start codon (Martí-Gómez *et al*. 2026b). In contrast to results in the Smn1 dataset, Local Epistasis Regression is roughly tied as the best performing model together with Variance Component Regression, although the Variance Component Regression model performs better for very small amounts of training data (Figure 3H).

Finally, we applied Local Epistasis Regression to estimate the structure of genetic interactions across sites in two protein fitness landscapes: (i) a complete combinatorial dataset in which nearly all possible combinations of amino acids at four positions in the binding domain of protein G were measured (GB1) (Wu et al. 2016), and (ii) a combinatorial landscape in which combinations of five amino acids at seven positions in the core of the SH3 domain of the FYN tyrosine kinase were quantified (FYN-SH3) (Escobedo *et al*. 2025). These datasets also show heterogeneous fitness correlations between pairs of sequences depending on the specific combination of sites at which they differ, which can be effectively captured by our estimated prior distribution (Figure S1A,B,D,E). The inferred parameters suggest that epistatic interactions are concentrated within specific subsets of sites (Figure S1C,F), consistent with patterns observed in the empirical RNA landscapes described above. For GB1, Local Epistasis Regression and Variance Component Regression are the best performing models and perform essentially identically over the whole range of training data sizes (Figure S1D), whereas for the SH3 domain Variance Component Regression performed better across the whole range of training data set sizes (Figure S1H). Thus, in summary, our analysis shows that empirical fitness landscapes often display heterogeneous correlations between pairs of sequences differing by the same number of mutations depending on the specific sites that are mutated, however, the extent to which incorporating this information into our prior distribution improves predictive performance varies across datasets relative to previous methods.

### The fitness landscape of a self-splicing intron

After validating the ability of Local Epistasis Regression to characterize the patterns of genetic interactions between pairs of sites in well-characterized fitness landscapes, we set out to study an 8-nucleotide fitness landscape of a self-splicing type I intron from *Tetrahymena thermophila* (Soo et al. 2021) that we have not previously analyzed. In particular, these 8 mutagenized nucleotides can form an extension of the P1 helix that is involved in recognition of the 5′splice site (Figure 4A). This P1 helix must then dissociate to form an alternative P10 helix with the 3′ splice site for catalyzing the splicing reaction between the 5′ and 3′ splice sites (Figure 4A). Thus, we expect a particularly rich structure of genetic interactions across sites, given the heterogeneous structural roles played by these nucleotides at different stages of the splicing reaction.

**Figure 4.**
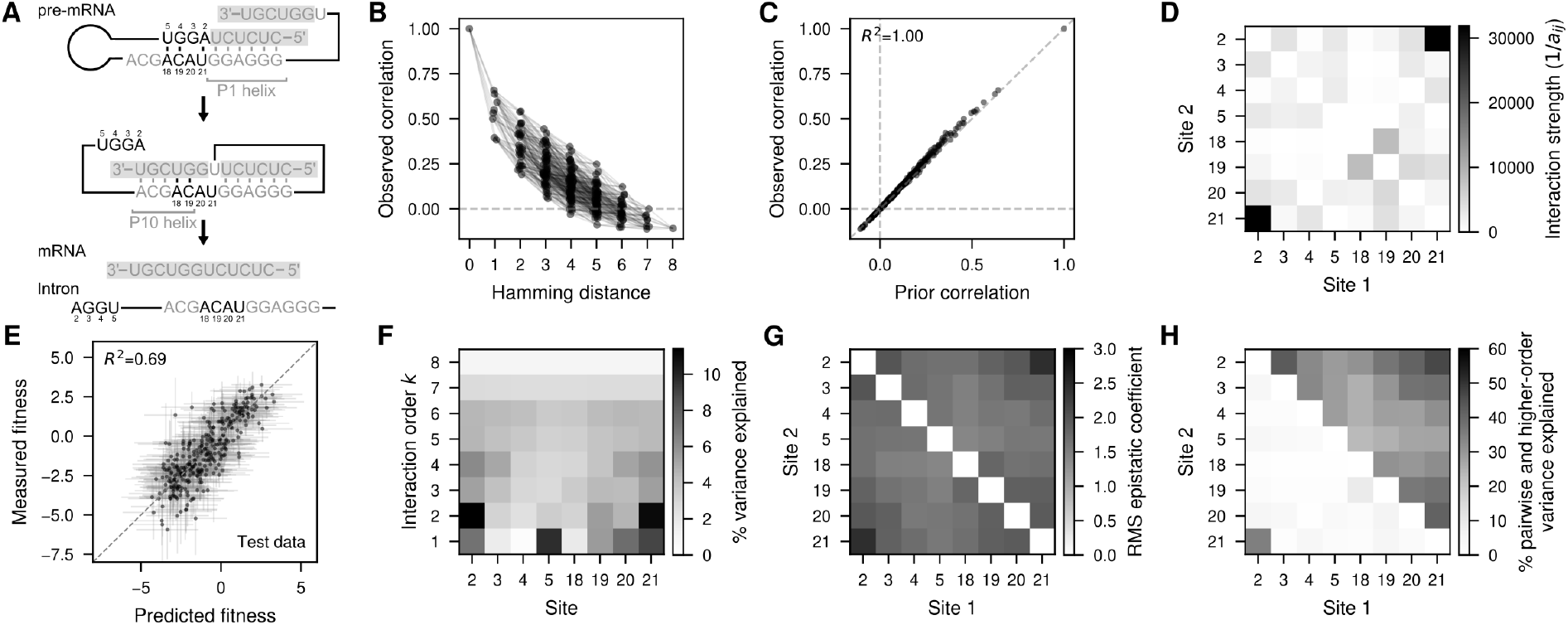
Inference and summary statistics of the fitness landscape of a self-spliced intron. (A) Schematic representation of the molecular mechanism of self-splicing in the model intron from *Tetrahymena thermophila* (Soo et al. 2021). Positions considered in the fitness landscape are highlighted in black and are numbered by their relative position in the intron. Other relevant sequences that are fixed in the background are shown in grey. Exonic sequences are shown with gray background. (B) Correlation in the measured fitness values for pairs of sequences differing at each possible subset of sites *D* arranged according to the Hamming distance *d* = |*D*|. Each dot represents a single distance class *D* and are joined by lines whenever the distance classes differ by a single position from each other. (C) Comparison of the observed correlation values in the data and the values under the estimated prior using Local Epistasis Regression. One correlation is shown for each possible set of positions *D* where two sequences may be differ (each dot represents a different *D*). (D) Heatmap representing the inferred model hyperparameters as 1/*a*_*ij*_ for every pair of sites *i, j* highlighting the patterns of genetic interactions across sites under the prior. (E) Measured values for held-out test sequences versus Local Epistasis Regression predictions. Horizontal error bars represent the 95% credible interval, whereas vertical error bars correspond to the 95% confidence interval under each measurement’s variance. (F) Heatmap representing the percentage of variance in the maximum a posteriori Local Epistasis Regression reconstruction explained by interactions of order *k* involving each position. (G) Root mean squared local double-mutant epistatic coefficient magnitude between mutations at each possible pair of predictions for the maximum a posteriori reconstruction. The plot indicates that relatively large local epistatic coefficients occur between mutations at essentially all pairs of positions. (H) Heatmap representing the percentage of variance in the maximum a posteriori reconstruction explained by pairwise (lower triangle) and higher-order (upper triangle) interactions that is explained by interactions involving pairs of positions.

We start again by computing the correlation in the measured fitness values for pairs of sequences differing at each possible combination of sites. The correlations, as before, depend not only on the number of sites at which two sequence differ, but more specifically on the combination of sites at which they do so, suggesting that different mutations have different effects on the predictability of other mutations (Figure 4B). We next estimate the parameters of the Local Epistasis Regression prior and find that the learned prior can again accurately recapitulate the observed correlations in the data (Figure 4C). The estimated *a*_*ij*_ values indicate that epistatic coefficients are not identically distributed across pairs of mutations at different combinations of sites, but that pairs of mutations at different sites tend to interact more strongly with each other (Figure 4D). The strongest signal comes from mutations at sites 2 and 21 which are by far the most strongly interacting positions, consistent with the P1 helix extension mechanism wherein position 2 basepairs with position 21 (Figure 4A).

Next, we perform inference of the complete fitness landscape under the learned prior while taking into account the estimated experimental error. These estimates accurately recapitulate the measured fitness in 0.5% (327) random held-out sequences (Figure 4E). Importantly, our Gaussian process formulation allows us to obtain uncertainty estimates for the fitness values of the held-out sequences showing good coverage properties, as the 95% posterior credible interval included the measured fitness for 91.1% of the held-out sequences. We found that all the regression models allowing higher-order interactions made similar predictions and had similar predictive performance (Figure S2A), and that in particular Local Epistasis Regression and Variance Component Regression both produced very similar fits (Figure S2B) and predictions for the set of held-out sequences (Figure S2C).

Next, we further explore the structure of epistasis in this dataset by computing a series of informative summary statistics from the maximum a posteriori (MAP) reconstruction of the complete landscape. First, we compute the percentage of variance explained by epistatic interactions of every possible order (Zhou *et al*. 2022) and find that the reconstructed landscape is highly epistatic, with only 42.3% of the variance explained by the additive component, 22.2% by pairwise interactions, and the remaining 35.4% explained by higher-order genetic interactions. To investigate how much each site contributes to interactions of different orders, we compute the percentage of variance explained by genetic interactions of order *k* that involve each site *i* (Figure 4F) (Martí-Gómez *et al*. 2026b). These statistics reveal substantial heterogeneity across sites. Some sites contribute little across all interaction orders, such as sites 4 and 18. Other sites contribute more strongly through their additive effects, such as position 5, whereas sites like 2 and 21 contribute primarily through pairwise and higher-order interactions. Next, we characterize the structure of genetic interactions between sites by computing the root mean square local double-mutant epistatic coefficient 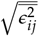 for every pair of sites (Figure 4G). These statistics reveal widespread epistatic interactions across pairs of sites, particularly among sites 2, 3, 20 and 21, and within the groups of positions 2–5 and 18–21, consistent with the regularization parameters (Figure 4D). However, as shown by Eq. 2, these quantities reflect contributions from both pairwise and higher-order interactions. To disentangle these contributions, we compute the percentage of pairwise 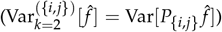 and higher-order variance 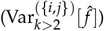 explained by interactions involving each pair of sites (Figure 4H). This analysis shows that a large fraction of pairwise epistatic variance is explained by interactions between positions 2 and 21 alone (Figure 4A, lower triangular part), whereas higher-order epistatic variance is more broadly distributed across sites, indicating that local double mutant epistatic coefficients for essentially all pairs of sites vary substantially across genetic backgrounds (Eq.3).

### Structure of the self-spliced intron fitness landscape

In order to better understand the qualitative structure of this fitness landscape, we apply a visualization method for producing low-dimensional representations of fitness landscapes where distances between genotypes in the representation reflect the expected waiting time to evolve from one genotype to another under a model of molecular evolution that includes mutation, selection, and drift (McCandlish 2011). Applying this method as implemented in the software package *gpmap-tools* resulted in Figure 5A (see Methods, Martí-Gómez *et al*. (2026b)). In the previous section we noted a particularly strong pattern of both pairwise and higher-order interactions involving positions 2 and 21 (Figure 4F), and we see that the visualization in Figure 5A largely separates sequences based on the nucleotides present at this pair of positions. Because the axes in such visualizations tend to highlight key barriers that make it difficult for a population to diffuse from one area of sequence space to another, we examined the mean fitness conferred by different combinations of alleles at these two sites for potential allelic incompatibilities (Figure 5B), revealing a consistent pattern wherein having both positions 2 and 21 occupied by pyrimidines (or also *C*_2_ *A*_21_) results in low mean fitness. This observation suggests that the P1 helix extension by base-pairing of positions 2 and 21 is not strictly necessary for functionality.

**Figure 5.**
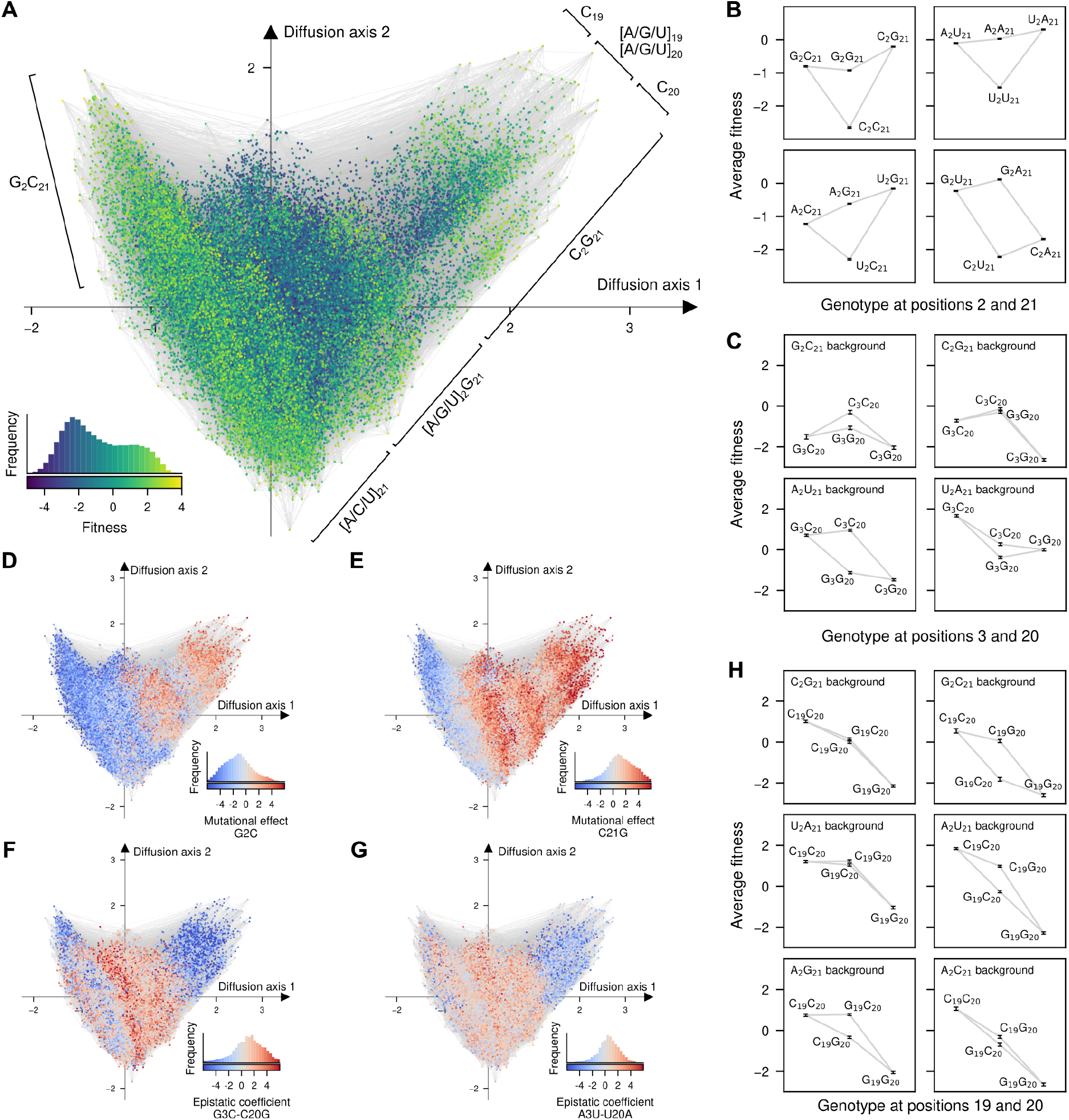
Visualization and characterization of the fitness landscape of a self-spliced intron. (A) Visualization of the inferred fitness landscape using Local Epistasis Regression. Every dot represents one of the possible 4^8^ possible sequences and is colored according to the predicted fitness. The inset represents the phenotypic distribution along with their corresponding color in the map. Sequences are laid out according to the first two Diffusion axes and dots are plotted in order according to Diffusion axis 3. (B,C,H) Diagrams representing average fitness for different subsets of sequences at positions 2 and 21 (B), for combinations of G and C alleles at positions 3 and 20 across different genetic contexts at positions 2 and 21 (C) and for combinations of G and C alleles at positions 19 and 20 across different genetic contexts at positions 2 and 21 (H). Indicated subsets of sequences are arranged along the x-axis according to Hamming distance from the leftmost subset. Error bars represent the 95% posterior credible intervals for these average fitness values. (D-G) Visualization of the inferred fitness landscape, as shown in (A), where nodes are colored by the mutational effect of G2C (D) and C21G (E), and the local epistatic coefficients between mutations G3C and C20G (F) and between A3U and U20A (G) when introduced in every possible genetic background throughout the landscape. The inset represents the distribution of the specific mutational effects or epistatic coefficients along with their corresponding color in the map.

Because the visualizations tend to spread apart high-fitness sequences that are separated by these types of incompatibilities, they are also useful for understanding how mutational effects and local epistatic coefficients vary across sequence space. Figures 5D and E show how *G*2*C* and *C*21*G* mutations are deleterious in the *G*_2_*C*_21_ cluster on the left-hand (negative) side of Diffusion Axis 1, but the same mutations become advantageous as one moves to the right-hand (positive) side. In fact, looking across all 8 positions, Figure S4 shows a widespread tendency for allelic preferences to change in a coherent manner across different clusters of sequences. Similarly, we can see how local double-mutant epistatic coefficients change in different regions of sequence space. For example, in the *C*_2_ *G*_21_ cluster on the right-hand side of Figure 5A, *G*3*C* and *C*20*G* are strongly negatively epistatic (Figure 5F) whereas in many other backgrounds such as *U*_2_ *A*_21_ background (Figure 5C), their interaction is strongly positive. Likewise the local double-mutant epistatic coefficient between *A*3*U* and *U*20*A* are strongly negative in the *C*_2_ *G*_21_ cluster but typically neutral or positive elsewhere (Figure 5G). Note that Figures 5F and 5G provide a simple illustration of the geometry of a 4th-order interaction, as we can see that the epistatic interactions at position 3 and 20 differ across the clusters of sequences defined by the base identities at positions 2 and 21.

The visualization also separates sequences within the *C*_2_ *G*_21_ background into three clusters, depending on the presence of *C* at positions 19 and 20 (Figure 5A). Moreover, Figure S4 shows that while *C*_19_ and *C*_20_ are generally preferred across the landscape, consistent with base pairing with cognate bases in the P10 helix in the second step of the splicing reaction (Figure 4A), the effects of alleles *C* and *G* at these two positions are strongly dependent on genetic background. To better understand these dependencies, we examined the average fitness of the four combinations of *C* and *G* alleles at positions 19 and 20 across different genetic contexts (Figure 5H). We find that in some genetic contexts, like *C*_2_ *G*_21_, *U*_2_ *A*_21_, and *C*_2_ *A*_21_, the base-pair–breaking mutations *C*19*G* and *C*20*G* are on average neutral or only mildly deleterious individually, but become substantially more deleterious when combined together. In contrast, in other genetic contexts, such as *G*_2_*C*_21_, *A*_2_*U*_21_, and *A*_2_*C*_21_, these mutations are more strongly deleterious individually and combine more additively. These epistatic interactions limit the accessible mutational paths between *C*_19_ *G*_20_ and *G*_19_*C*_20_ sequences, and explain why these clusters appear separated in the visualization (Figure 5A). The background dependence of this interaction provides an additional illustration of how higher-order interactions in this system can be largely understood in terms of the identities at positions 2 and 21, and how the alleles at these two positions modulate epistatic interactions between the other sites.

## Discussion

In this work, we propose a new method for inferring empirical fitness landscapes from experimental data. Our method is based on the idea that genetic interactions are not uniformly distributed across sites; rather, some sites tend to interact more strongly with specific other sites. This information about sets of sites exhibiting stronger epistatic interactions can be extracted from how correlations between measured fitness values depend on the specific combinations of sites at which sequences differ. We then incorporate this information into a prior distribution defined over all possible fitness landscapes for Bayesian inference of complete combinatorial landscapes from incomplete and noisy data. Importantly, although our prior allows the incorporation of different degrees of regularization for epistatic coefficients between different pairs of sites, it still allows these coefficients to vary across genetic backgrounds, thereby preserving the benefits of other methods that contain genetic interactions of all orders (Zhou and McCandlish 2020; Zhou *et al*. 2022). Applying this method to three experimental datasets revealed that mutations do tend to interact more strongly with other specific mutations in empirical landscapes (Figures 3C,F, S1C,G and 4D), often reflecting known molecular interactions between positions (Figures 4A,D). Finally, we performed an in-depth analysis of the inferred landscape for a self-splicing intron, where we found that the nucleotide identities at positions 2 and 21, which can potentially form an extension of the P1 helix, determine the nature of genetic interactions at many other pairs of positions. This shows how higher-order interactions, which may at first seem mysterious, can be understood in terms of local pairwise interactions that change coherently as one moves from one region of the fitness landscape to another.

Our method relies on a simple summary statistic that quantifies the overall strength of local genetic interactions between a given pair of mutations across across all genetic backgrounds. Specifically, we compute the average squared epistatic coefficient between mutations at every possible set of sites. While this statistic was previously derived in the context of the *γ*_*i* →*j*_ measure (Ferretti *et al*. 2016), here we show that it can be expressed as a quadratic form with a positive semi-definite matrix that admits a Kronecker product decomposition into *ℓ* smaller matrices. This allowed us to derive (i) the relationship between the total epistatic variance of all orders involving any pair of sites *i* and *j* (Reddy and Desai 2021; Martí-Gómez *et al*. 2026b) and the average squared size of local double-mutant epistatic coefficients defined by pairs of mutations at sites *i* and *j*, and (ii) the relationship between the higher order epistatic variance and the variance in these epistatic coefficients across genetic backgrounds, clarifying how local epistatic effects and their context dependence connect to the global variance decomposition of fitness landscapes. Interestingly, while the ability to conduct this Kronecker factorization enables efficient computation of these summary statistics in combinatorially complete fitness landscapes (Martí-Gómez *et al*. 2026b,a), it also opens up the possibility of calculating these statistics for astronomically large fitness landscapes for the maximum a posteriori estimate under a certain class of Gaussian process models (Petti *et al*. 2025), which would allow the incorporation of higher-order interactions in applications such as protein contact prediction (Marks *et al*. 2012) and 3D-structure inference from mutagenesis experiments (Schmiedel and Lehner 2019). Finally, the Kronecker 10 Local epistasis regression factorization form makes it straightforward to generalize these statistics in two directions: (i) computing squared epistatic coefficients between specific mutations within the same pair of sites (see Supplement) and (ii) averaging of the squared coefficients across different genetic backgrounds or over backgrounds drawn from site-factorizable probability distributions (rather than from the uniform distribution as we have done here).

As in previous work (Zhou and McCandlish 2020; Chen *et al*. 2021; Zhou et al. 2022; Reddy and Desai 2021; Zhou et al. 2025), we use our mathematical understanding of this summary statistic to define random field models of fitness landscapes, which can be used as prior distributions for Gaussian process inference of empirical landscapes from high-throughput experimental data. Our prior allows the same mutation to have different effects on the predictability of different mutations, whereas previous approaches assign all mutations (Zhou and McCandlish 2020; Chen et al. 2021) or each individual mutation (Reddy and Desai 2021; Zhou et al. 2025) a constant effect on the predictability of other mutations, preventing them from encoding structured interaction patterns between sites. Despite finding widespread evidence that specific pairs of sites tend to interact more than others, incorporating this realism into our prior did not always increase predictive performance. One potential reason for this is that while more flexibly encoding differences between pairs of sites, our model makes assumptions about the form of higher-order epistasis that may be more appropriate for some datasets than others. A second potential reason for this variable performance concerns our kernel alignment procedure for choosing the hyperparameters. We had previously seen a modest improvement in model performance for Empirical Variance Component Regression when the hyperparameters were chosen by evidence maximization rather than kernel alignment (Zhou et al. 2025), suggesting that the two modes of inference should generally have similar performance. However, here we see that the connectedness model fit by kernel alignment performed far worse on the GB1 and Smn1 datasets than our previous implementation using evidence maximization (Zhou et al. 2025), suggesting that the method of hyperparameter optimization might have a greater impact than previously thought. Unfortunately, unlike kernel alignment, which we were able to implement efficiently using the fact that the covariance under our model only depends on the subset of sites at which two sequences differ, implementing evidence maximization for Local Epistasis Regression as in Zhou *et al*. (2025) is more challenging due to the need to calculate and combine 2^*ℓ*^ kernels.

A fundamental aspect of fitness landscapes is their dependence on the environment. Increasingly, datasets measure fitness across multiple environmental conditions, necessitating models that allow mutational effects and epistatic interactions to vary across environments (Nguyen Ba et al. 2022; N’Guessan *et al*. 2025; Bakerlee et al. 2022; Soo et al. 2021; Ishigami et al. 2024). Our framework naturally extends to this setting by treating environmental conditions as additional loci, enabling the inference of priors in which mutational effects depend jointly on genetic background and environment. Importantly, our prior can learn and encode that mutations tend to change by different magnitudes when introducing an additional mutation compared to when changing environments, effectively allowing different prior variances for gene-by-gene and gene-by-environment interactions.

Our method shares several limitations with previous Gaussian process approaches (Zhou and McCandlish 2020; Zhou *et al*. 2022, 2025). First, it does not explicitly model non-specific or global epistasis (Otwinowski *et al*. 2018; Domingo *et al*. 2019), but instead relies on specific epistatic interactions to fit these global dependencies, potentially limiting the interpretability of the estimated hyperparameters as the structure of specific genetic interactions between sites. Second, by taking an empirical Bayes approach, in which we first estimate the parameters of the prior and then use those parameters for inference of the fitness landscape, we are not taking into account potential uncertainty in the estimation of these parameters. Our new model may be more sensitive to this limitation than previous approaches because of the larger number of hyperparameters that need to be estimated (Zhou and McCandlish 2020; Zhou *et al*. 2022). Third, our current implementation takes advantage of the mathematical structure of the covariance matrix to allow efficient computation of the posterior distribution (Equations 7 and 8) without explicitly building the covariance matrix for complete fitness landscapes. While this trick allows us to compute the posterior distribution for fitness landscapes containing hundreds of thousands of sequences, it is limited to sequences of relatively short length (about 5 amino acids, 12 nucleotides or 24 bi-allelic loci). Moreover, the need to evaluate 2^*ℓ*^ different kernels, even if the number of hyperparameters is much lower, hinders the applicability even under GPU-accelerated frameworks for scalable Gaussian process inference (Gardner *et al*. 2018; Wang *et al*. 2019) that facilitated the application of previous models to fitness landscapes defined over longer sequences (Zhou *et al*. 2025).

## Materials and methods

### Fitness landscape of a 5′ splice site sequence

Data reported by Wong *et al*. (2018) was processed as previously reported (Zhou *et al*. 2022, 2025). Briefly, we assumed a log-normal distribution of enrichment ratios across 1-7 replicates, for each different 5′ splice site sequence *x*. The bias corrected geometric mean of the enrichment ratio was used as an estimate of the median enrichment ratio when the enrichment ratio was strictly positive for all replicates. Otherwise, the median of enrichment scores was used to estimate the phenotype *y*_*x*_. Sequence-specific variance *y*_*x,var*_ was estimated as indicated below, where 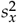 is the sample variance of the log-enrichment ratios if all replicates were strictly positive and were measured in at least two samples or the median of all 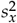 for sequences *x* with at least two replicate measurements:

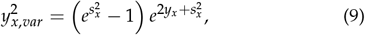

see (Zhou *et al*. 2022) for more details.

### Fitness landscape of the Shine-Dalgarno sequence

Data reported by Kuo *et al*. (2020) was processed as previously reported (Martí-Gómez *et al*. 2026b). Briefly, fitness was estimated as the mean log(GFP) for 257,565 measured sequences with a common measurement variance of *s*^2^ = 0.058 using genotypes measured across all three experimental replicates. The squared standard error for each genotype *i* was computed by dividing this observed experimental variance *s*^2^ by the number of replicates *n*_*x*_ in which each sequence was measured (*y*_*x,var*_ = *s*^2^/*n*_*x*_).

### Fitness landscape of protein G binding domain

Data was processed as previously described (Zhou and McCandlish 2020; Zhou *et al*. 2022). Briefly, we used the number of sequencing reads for each sequence *x* in the input sample 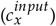 and in the selected sample 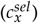 reported in (Wu et al. 2016) to estimate the log-enrichment ratio relative to the wild-type sequence *y*_*x*_ as a measure of the binding strength. Moreover, we estimated the error variance *y*_*x,var*_ of this estimate (Rubin *et al*. 2017):

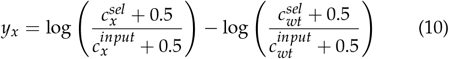

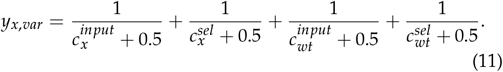

### Fitness landscape of the FYN protein SH3 domain

Data was used as reported by the original study (Escobedo *et al*. 2025). Specifically, we downloaded the processed data from GEO (GSE266299) used the scaled relative fitness measurements and the reported experimental errors for our downstream analysis.

### Fitness landscape of a self-splicing intron

The number of reads for each variant across six-replicates in the presence and absence of Kanamycin was collected at 30º from the original study (Soo et al. 2021). Following Soo *et al*. (2021), fitness *y*_*x*_ for each sequence *x* was estimated as the log_2_(Fold change) between Kanamycin treated and control samples using PyDESeq2 (Muzellec et al. 2023) without shrinkage towards zero, and squared standard errors were kept as measure of experimental variance *y*_*x,var*_ for downstream analysis. Local Epistasis Regression was used to estimate the complete combinatorial fitness landscape taking into account the experimental errors for each of the sequence using all available data except 0.5% (327) sequences. We computed the posterior mean and variance for these 327 sequences to evaluate the performance of the model in unobserved sequences. Using the posterior mean for the complete fitness landscape, we computed the variance explained by epistatic interactions of every possible order for each site and between pair of sites using *gpmap-tools* (Martí-Gómez et al. 2026b). We then generated a low-dimensional representation in which distances between pairs of sequences reflect the expected time to evolve from one sequence to the other (McCandlish 2011) under an evolutionary model in the weak mutation regime as implemented in *gpmap-tools* (Martí-Gómez et al. 2026b). We generated visualization coordinates under different strengths of selection by choosing different values for the expected fitness under longterm mutation-selection-drift (i.e. expected fitness at stationarity, Figure S3). A long-term expected fitness of 1.6 (corresponding to the 87% percentile in the distribution of fitness values) was used for the final visualization.

## Data and code availability

The methods presented in this work have been implemented in *gpmap-tools* v0.4.2 (Martí-Gómez et al. 2026b), an open-source library available at https://github.com/cmarti/gpmap-tools. Code and data to reproduce the analyses presented in this paper are available at https://github.com/cmarti/deltaU.

## Funding

CMG and DMM were supported by the US National Institutes of Health (NIH) grant R35GM133613 and additional funding from the Simons Center for Quantitative Biology at Cold Spring Harbor Laboratory. This work was performed with assistance from the NIH Grant S10OD028632.

## Conflicts of interest

The authors declare no conflicts of interest.

## Supplementary Information

### Epistasis on fitness landscapes

Let *f* be an *α*^*ℓ*^-dimensional vector encoding the fitness associated with each genotype in the space of possible haploid sequences with *α* alleles and *ℓ* sites *S* = {1, 2, …, *ℓ*}. In this section, we review two common ways to quantify epistasis in a fitness landscape: one based on local epistatic coefficients (Zhou and McCandlish 2020; Chen et al. 2021) and a second one based on the variance explained by genetic interactions of different orders (Stadler and Happel 1999; Zhou et al. 2022) or across subsets of sites (Martí-Gómez et al. 2026b).

#### Variance components

Any fitness landscape *f* can be decomposed into orthogonal components *f*_*k*_ corresponding to genetic interactions of order *k*

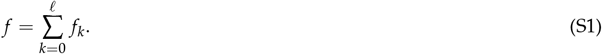

These components can be obtained by projecting the *f* into the *k*th-subspace using the orthogonal projection matrix *P*_*k*_ given by the Krawtchouk polynomials (Stadler and Happel 1999; Zhou et al. 2022):

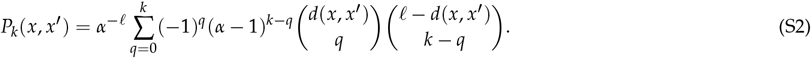

Moreover, each *k*-th order subspace can be further decomposed into orthogonal components corresponding to the contributions of interactions between specific subsets of sites *U* ⊆ *S* of size *k*:

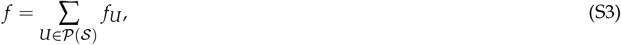

where *P*(*S*) is the set of all subsets of *S* (i.e. the power set of *S*) and *f*_*U*_ = *P*_*U*_ *f*, where *P*_*U*_ is an orthogonal projection matrix onto the corresponding subspace. Each such *P*_*U*_ can be expressed as a Kronecker product of a series of site-specific projection matrices onto the constant subspace, defined by 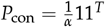, and the orthogonal or additive subspace, given by *P*_add_ = *I* − *P*_con_. In particular

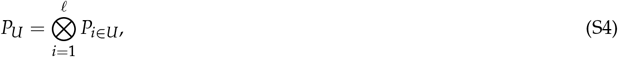

where *P*_*i*∈*U*_ = *P*_add_ when site *i* ∈ *U* and *P*_*i*∈*U*_ = *P*_con_ otherwise (Martí-Gómez et al. 2026b):

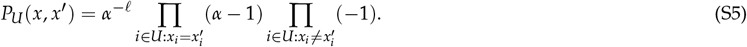

Because *U*-components are orthogonal to each other, the total variance in a fitness landscape can be expressed as a sum of variances explained by genetic interactions of order *k* (i.e. summing over all *U* with |*U*| = *k*) or by interactions between the sites *U*, respectively (Martí-Gómez et al. 2026b):

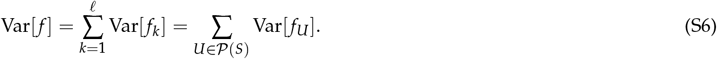

#### Local epistatic coefficients

The classical way to quantify epistasis is through the definition of the epistatic coefficient *ϵ* for a pair of mutations at different loci *A* → *a* and *B* → *b*, which quantifies the difference in the effect of mutation *A* → *a* in the presence of allele *B* compared to that in the presence of allele *b* at the other locus:

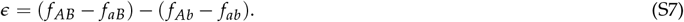

This epistatic coefficient is defined locally, as it depends only on four specific genotypes sharing the same genetic background. Still, we can compute the average magnitude of local epistatic interactions 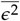 in a given fitness landscape by averaging the its squared values across every possible pair of mutations at every possible genetic background:

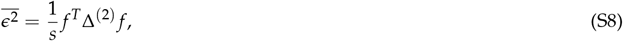

where 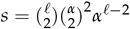 is the number of epistatic coefficients and Δ^(2)^ is a positive semi-definite sparse matrix (Zhou and McCandlish 2020; Chen et al. 2021). These results can be generalized to quantify local epistatic coefficients of any order *P* using

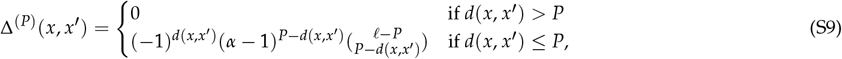

where *d*(*x, x*′) represents the Hamming distance i.e. number of single point mutations, separating sequences *x* and *x*′, such that the sum of the squared *P*-th order local epistatic coefficients is given by *f* ^*T*^ Δ^(2)^ *f* and the mean sqaured *P*-th order local epistatic coefficient is obtained by dividing by the number of such coefficients 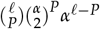 (Chen et al. 2021). Interestingly, Δ^(*P*)^ can also be expressed as a weighted sum of the projection operators into the *k*-th order subspaces given by *P*_*k*_ :

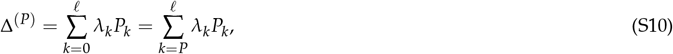

where 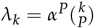 (Chen et al. 2021) correspond to the eigenvalues of the Δ^(*P*)^ operator. Note that the second equality arises because 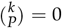 for *k < P*, corresponding to the fact that all local *P*-th order epistatic coefficients are zero for any function *f* with maximal order less than *P* (so that all such functions are in the null space of Δ^(*P*)^) and that moreover alteration of any component of order less than *P* for an arbitrary *f* leaves all *P*-th order local epistatic coefficients unchanged.

### Epistatic coefficients among subsets of sites

In this section, we describe how the Δ^(*P*)^ operator that extracts the sum of squared epistatic interactions in a fitness landscape can be decomposed into the sum of simpler operators that we call Δ^(*U*)^, corresponding to local epistatic interactions only among a subset of sites *U* ⊆ *S*. These results generalize the main text results focused on local epistatic interactions between pairs of sites *i* and *j* i.e. *U* = {*i, j*}.

Let *E*_*U*_ be a *s*_*U*_ × *α*^*ℓ*^ matrix such that the entries in the *E*_*U*_ *f* vector encode all the local |*U*|-th order epistatic coefficient for a subset of positions *U. s*_*U*_ is the number of different epistatic coefficients corresponding to local |*U*|-th order mutant epistatic coefficients within sites *U*, and is given by the product of the number of genetic backgrounds at sites not in *U α*^*ℓ*−|*U*|^ and the number of possible combinations of mutations across the sites 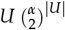:

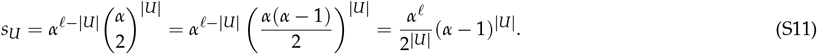

Thus,

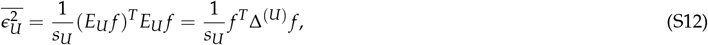

where the entries of Δ^(*U*)^ for a pair of sequences *x* and *x*′ can be obtained by summing over all possible local |*U*|-mutant epistatic coefficients between sites in *U*

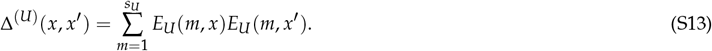

*E*_*U*_ (*m, x*′) = 0 if sequence *x*′ is not involved in epistatic coefficient *m*, and takes values −1 or 1 otherwise. Thus, we only need to sum overlocal epistatic coefficients involving both *x* and *x*′. If 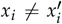 for any position *i* ∈/*U*, then sequences *x* and *x*′ cannot be involved in any epistatic coefficient and thus Δ^(*U*)^(*x, x*′) = 0. Otherwise, *E*_*U*_ (*m, x*)*E*_*U*_ (*m, x*′) = (−1)^*d*(*x,x*′)^ depending on the Hamming distance between them *d*(*x, x*′). Moreover, the number of local |*U*|-mutant epistatic coefficients that involve both s*x* and *x*′ is given by (*α* ™ 1) ^|*U*|™d(x,x ′)^. Thus

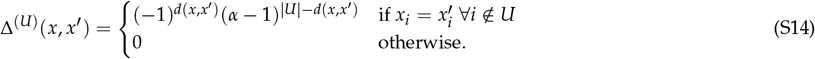

We can verify that we can recover the Δ^(*P*)^ operator by summing Δ^(*U*)^ over all possible subsets of sites *U* of size *P*. If *d*(*x, x*′) *> P*, there is no single set of sites *U* for which the context at sequences *x* and *x*′ can match and thus ∑_*U*:|*U*|=*P*_ Δ^(*U*)^(*x, x*′) = 0. For *d*(*x, x*′) ≤ *P*, the entries of Δ^(*U*)^(*x, x*′) are the same but they are summed over multiple *U*. As we only sum over *U*’s such that *x* and *x*′ share the context, the number of times we are summing them corresponds to the number of ways we can choose *P* − *d*(*x, x*′) sites that match out of the *ℓ* − *P* sites in the shared context between them. Thus,

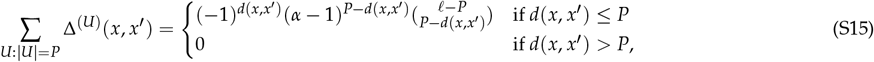

which exactly matches the Δ^(*P*)^ operator (Zhou and McCandlish 2020; Chen et al. 2021).

### Relationship between local epistatic coefficients and variance components

In this section, we describe some properties of the new Δ^(*U*)^ operator and the relationships with the projection operators into the subspaces corresponding to interactions of different orders and subsets of sites *U*.

One useful property of the Δ^(*U*)^ operator is that it can be expressed as a Kronecker product of site-specific matrices

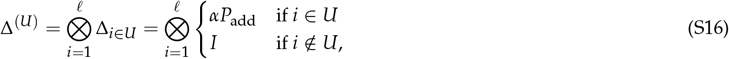

with entry-wise formula given by:

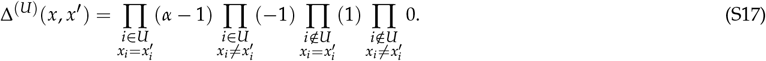

Thus, we can use the mixed-product property to show that the columns of *P*_*U*_′ are eigenvectors of Δ^(*U*)^ with eigenvalue *α*^|*U*|^ if *U* ⊆ *U*′ or are in the null space of Δ^(*U*)^ whenever *U* ⊈ *U*′.

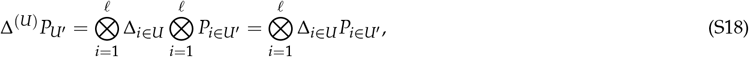

where

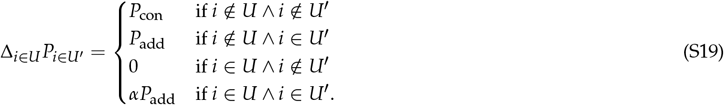

Therefore

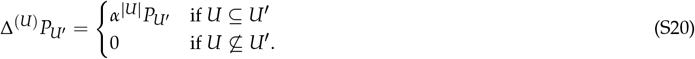

Because the projection operators *P*_*U*_′ are orthogonal to each other, we can express Δ^(*U*)^ as a linear combination of *P*_*U*_′ weighted by their eigenvalues (either 0 or *α*^|*U*|^):

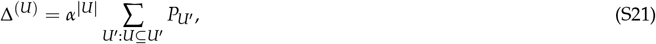

so that, using the fact that sums of projection matrices into orthogonal subspaces are themselves projection matrices, we see that Δ^(*U*)^ is itself just a |*U*|-dependent constant times a specific projection matrix. Moreover, this relation between Δ^(*U*)^ and the *P*_*U*_′ with *U* ⊆ *U*′ can be used to derive the relationship between the Δ^(*P*)^ operator and the projection operator into the *k*th order subspace *P*_*k*_ :

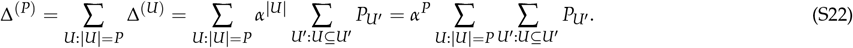

It is now easy to see that the number of times we are summing each *P*_*U*_′ depends on the size of *U*′. In particular, it corresponds to the number of *U* : |*U*| = *P* subsets in *U*′, which can be calculated as the number of ways of choosing |*U*| = *P* sites out of |*U*′ |. Then, we can use the fact that *P*_*k*_ = ∑_*U*: *U* =*k*_ *P*_*U*_ (Martí-Gómez et al. 2026b) to recover the eigendecomposition of the Δ^(*P*)^ operator (Chen *et al*. 2021):

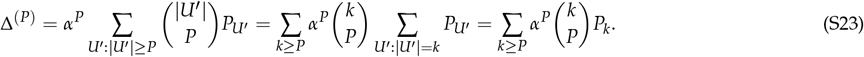

### Relationship between local |*U*| **-way epistatic coefficients and the epistatic variance explained by the subset** *U*

A useful low dimensional summary statistic to describe the patterns of genetic interactions across subsets of sites in a fitness landscape is the epistatic variance explained by a set of sites *U* (Crawford et al. 2017; Reddy and Desai 2021; Martí-Gómez et al. 2026b). We define this variance Var^(*U*)^[*f*] to include not only the variance explained by |*U*|-way interactions between mutations at the sites |*U*|, but also all other interactions of order higher than |*U*| where all sites in *U* are involved, so that Var^(*U*)^[*f*] provides an overall measure of the amount of |*U*|-way and higher epistasis that the subset of sites *U* is involved in. This quantity can be computed by projecting the fitness landscape *f* onto the 2^*ℓ*^ subspaces defined by every possible *U*′ given by *f*_*U*_′ = *P*_*U*_′ *f*, summing over the *f*_*U*_′ ‘s of interest and computing the inner product.

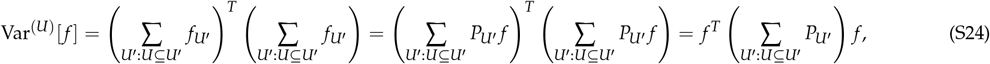

Where

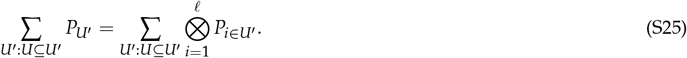

As the Kronecker product is commutative up to row and column permutations given by *Q*_1_ and *Q*_2_, we can write the factors in order depending on whether the sites are in *U* or not:

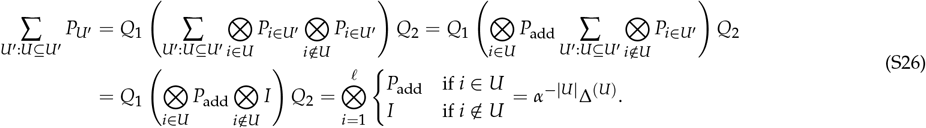

Thus, this shows that the sum of squared epistatic coefficients among a subset of sites *U* given by *f* ^*T*^ Δ^(*U*)^ *f* is equal to the epistatic variance explained by those sites Var^(*U*)^[*f*] multiplied by a factor of *α*^|*U*|^. Noting that 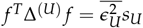 and simplifying *s*_*U*_ /*α*^|*U*|^ yields Equation 2 in the main text. Similarly, the sum of squared local *P*-epistatic coefficients across the complete fitness landscape given by *f* ^*T*^ Δ^(*P*)^ *f* can also be interpreted as the sum the epistatic variances of all possible combinations of *P* sites multiplied by a factor of *α*^*P*^.

### Relationship between mean and variance of local |*U*|-way epistatic coefficients and the variance components

In the previous sections, we have shown the relationships between the average squared epistatic coefficients between mutations at sites *U* and the total variance explained by interactions of order |*U*| or larger involving all sites in *U*. In this section, we describe how these quantities relate to the mean and variance of the distribution of |*U*|-way epistatic coefficients for a set of mutations within sites *U*. To do so, we first decompose the epistatic coefficients between mutations at sites *U* into the epistatic coefficients for each possible combination of mutations at sites *U*. Using this more granular statistic, we derive the mean of the epistatic coefficients between a given set of mutations, and use its square, together with the average squared epistatic coefficient to compute the variance. Finally, we take the average of these variances over all possible combinations of mutations at sites *U* and derive its relationship to the variance components defined over the subsets of sites *U*.

Let *C* = {{*c*_11_, *c*_12_},…., {*c*_*k*1_, *c*_*k*2_}} be the set of *k* pairs of characters at a set of *k* positions *U* and *ϵ*_*UC*_ be the *α*^*ℓ*−*k*^-dimensional vector of *k*-way epistatic coefficients between the mutations specified by *C* across every possible genetic background. Let *E*_*UC*_ be an *α*^*ℓ*−*k*^ × *α*^*ℓ*^ such that *ϵ*_*UC*_ = *E*_*UC*_ *f* and note that this matrix that can be expressed as a Kronecker product of site specific factors:

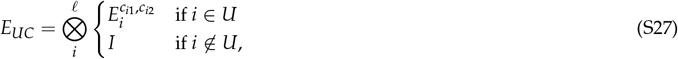

where 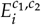 is a row vector with entries given by

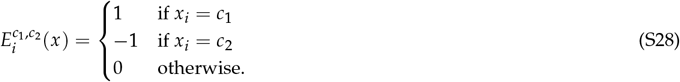

Using these expressions, we can compute the average squared epistatic coefficients restricted to the sets of mutations defined by *C* as

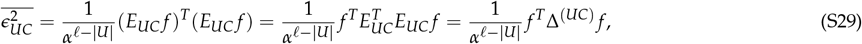

where Δ^(*UC*)^ can be easily computed by using the Kronecker factorization of *E*_*UC*_

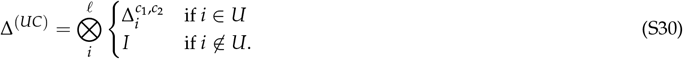

The factor 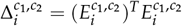 takes the form

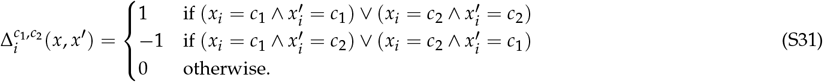

It is easy to see that 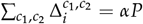 and that Δ^(*U*)^ can be expressed as a sum of Δ^(*UC*)^ for all possible combinations of pairs of alleles *C* at sites *U*.

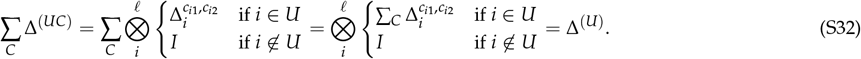

Next, we use the *E*_*UC*_ to compute the average epistatic coefficients 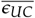 between the set of mutations defined by *C* at sites *U* across all possible genetic backgrounds:

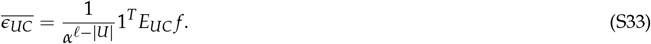

This quantity corresponds to the previously proposed background-averaged epistatic coefficient for parametrizing and describing sequence-function relationships (Faure et al. 2024a; Petti et al. 2025). As the sign of 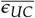 depends on the ordering of alleles *c*_*i*1_ and *c*_*i*2_ for each position *i*, we can use its squared value 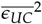 to describe the magnitude of the average epistatic coefficients independently of the choice of reference allele at each position:

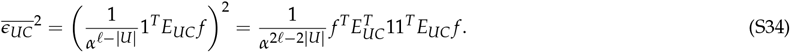

Let 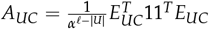, so that 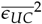 can be expressed as a function of a quadratic form with this matrix given by 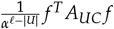. To derive a simple expression for *A*_*UC*_ and understand its relationship with other objects, we use the fact that all of the matrices *E*_*UC*_ and 11^*T*^ can be expressed as Kronecker products of site specific factors:

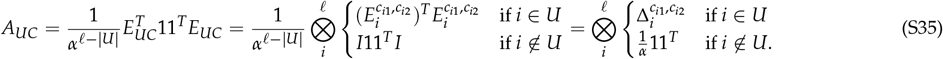

It is easy to see that *A*_*UC*_ is in the subspace defined by genetic interactions of order *U* between sites *U* since *P*_*U*_ *A*_*UC*_ = *A*_*UC*_. In fact, if we sum over all possible combinations of mutations *C*, we obtain a matrix that is proportional to the projection operator into the *U*-subspace *P*_*U*_ :

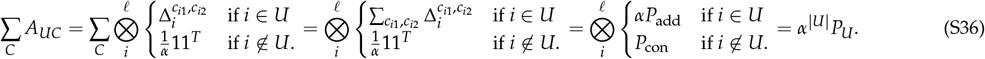

Using this equivalence, we can show that the average squared mean epistatic coefficient for mutations at sites *U* is proportional to the variance explained by genetic interactions of order |*U*| between those sites:

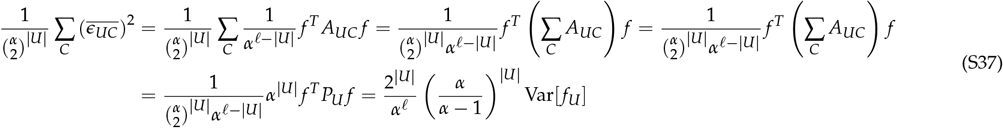

Finally, we can derive the variance over local epistatic coefficients between mutations at sites *U* by using the relationship between the variance and the raw second moments 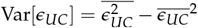. as well as the average variance across all possible sets of mutations *C* at sites *U* and show that it is proportional to the variance explained by genetic interactions of order higher than |*U*| involving sites *U*:

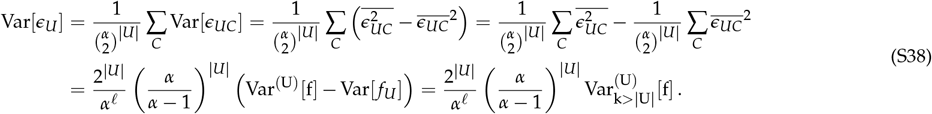

### Covariance function between sequences differing at subsets of sites

In addition to characterizing interaction structure through squared epistatic coefficients or epistatic variances of specific subsets of sites *U*, it is often useful to summarize an *arbitrary* landscape *f* by how similar fitness values are for sequences that differ at particular sets of sites. In particular, let *H*(*x, x*′) be the subset of sites at which two sequences *x* and *x*′ differ

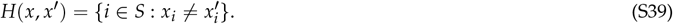

We define the covariance function for a mismatch set *D* as

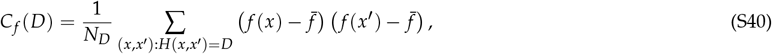

where 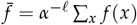 and *N*_*D*_ = *α*^*ℓ*^(*α* − 1)^|*D*|^ is the number of sequence pairs that differ at sites *D*.

*C*_*f*_ (*D*) can be expressed as a quadratic form 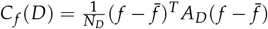, where *A*_*D*_ is an *α*^*ℓ*^ × *α*^*ℓ*^ matrix given by

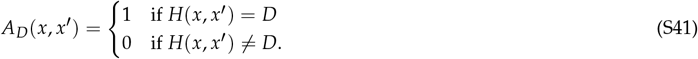

This matrix can be written as a Kronecker product of single-site matrices, enabling efficient computation of this quantity without explicit evaluation of the matrices *A*_*D*_ (Martí-Gómez et al. 2026b):

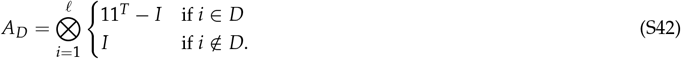

Moreover, if we consider the decomposition of *f* into its orthogonal components *f*_*U*_ = *P*_*U*_ *f* such that *f* = ∑_*U* ∈ *P* (*S*)_ *f*_*U*_, we can express the covariance function as a sum of covariance over all the possible *U*-components:

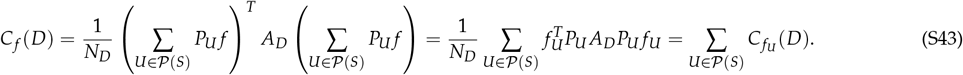

Next, we note that *P*_*U*_ *A*_*D*_*P*_*U*_ can be easily calculated through multiplication of their Kronecker factors individually:

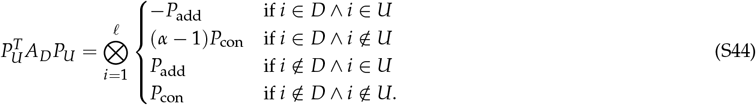

In fact, this can be summarized as 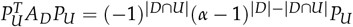. Since *f*_*U*_ is in the *U*-subspace, *P*_*U*_ *f*_*U*_ = *f*_*U*_ and thus

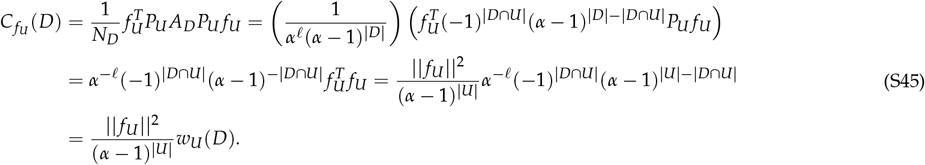

Here, we denote by *w*_*U*_ (*D*) the contribution of genetic interactions among sites *U* to the covariance between sequences that differ at sites *D*. We refer to these quantities as subset-resolved covariance weights and they are related to the classical Krawtchouk polynomials *w*_*k*_ (*d*) arising in the Fourier analysis of fitness landscapes (Stadler 1996; Zhou et al. 2022). In particular, the functions *w*_*k*_ (*d*), which depend only on the Hamming distance *d* = |*D*|, are recovered by summing *w*_*U*_ (*D*) over all subsets *U* of size *k*,

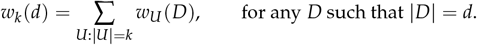

### Gaussian random field landscapes

In the previous sections, we have explained different ways to characterize the patterns of epistatic interactions in a given fitness landscape *f*. Here we aim to define a probabilistic ensemble of fitness landscapes via a Gaussian distribution p(*f*). This Gaussian distribution can be used as a theoretical random fitness landscape model, similar to the classical random fitness landscape models e.g. Rough Mount Fuji, NK or House of Cards landscapes (Kingman 1978; Kauffman and Weinberger 1989; Aita and Husimi 1998), or as a prior distribution for Gaussian process inference

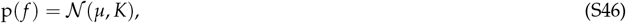

as previously proposed (Zhou et al. 2022, 2025). Here, we generally assume that the prior Gaussian distribution has zero mean i.e. *µ* = 0. To define this prior, let *Q*_*U*_ be an *α*^*ℓ*^ × (*α* − 1)^|*U*|^-dimensional matrix containing an orthonormal basis for the subspace defined by |*U*|-way interactions among the sites *U* (i.e. the column space of *P*_*U*_). We assume that the coefficients for these basis vectors are drawn independently from a zero-mean Gaussian with variance *λ*_*U*_. In particular, let *b*_*U*_ be the (*α* − 1)^|*U*|^-dimensional vector of coefficients:

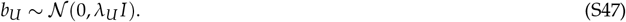

If we then define *f* = ∑_*U* (*S*)_ *Q*_*U*_*b*_*U*_, it is easy to see that *f* is also a zero-mean Gaussian distribution with covariance matrix *K* given by

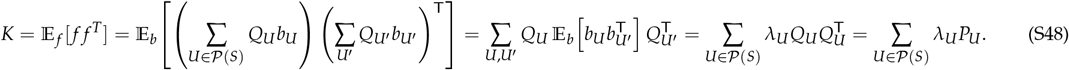

Since the columns of *Q*_*U*_ are orthonormal, *P*_*U*_ corresponds to the projection matrix into the function subspace spanned by interactions between sites in *U*, as given by Eq. S4. Here, we note that the entries of *P*_*U*_ (*x, x*′) only depend on the Hamming distance between sequences *x, x*′ at sites in *U*, but not on the alleles at which they match or differ. Thus, the covariance matrix *K*(*x, x*′) only depends on the subset of sites at which *x* and *x*′ differ.

Moreover, the variance of the regression coefficients *λ*_*U*_ for each subset of sites *U* determines the expected variance explained by interactions between exactly sites in *U* and is given by E _*f* ~N (0,*K*)_ || *f*_*U*_ ||^2^ = (*α* − 1)^|*U*|^ *λ*_*U*_.

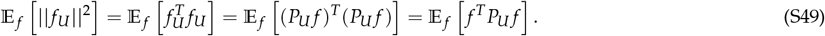

We note that samples from the random field model *f* ~ N (0, *K*) can be drawn by first drawing *z* ~ N (0, *I*) and then computing 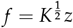, such that

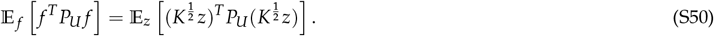

Moreover, as the columns of *P*_*U*_ are eigenvectors of *K* with eigenvalue *λ*_*U*_ and *P*_*U*_ matrices are orthogonal to each other, 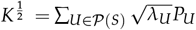 and thus

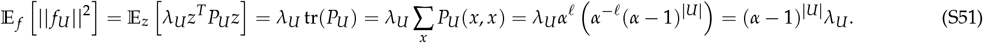

### A prior distribution for fitness landscapes

Because for sequences of length *ℓ* these priors are defined by there 2^*ℓ*^ values *λ*_*U*_, defining and interpreting this large number of the parameters can be challenging. Here, we use the relationship between the Δ^(*U*)^ and the projection operators into the *U*-subspace shown in Eq. S21 to define a simplified prior where the *λ*_*U*_ corresponding to interactions of order higher than *P* are parametrized via the different 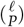 values of *a*_*U*_ :

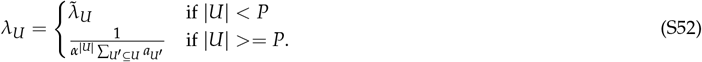

To understand the role of the parameters *a*_*U*_ in the prior, we can write it as

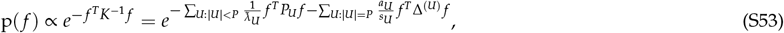

where *s*_*U*_ is the number of local epistatic coefficients between sites in *U* and the value of *a*_*U*_ modulates how much the prior penalizes these coefficients for a given *f*. Interestingly, one can see that as *a*_*U*_ → ∞, *λ*_*U*_ approaches 0 for all function subspaces explained by interactions involving the sites in *U*, essentially removing interactions of order equal or higher than |*U*| involving the set of sites *U*. For instance, for *P* = 2, setting *a*_*ij*_ = ∞ enforces the assumption that there are no genetic interactions between sites *i* and *j* or, equivalently, that the effects of mutations at site *i* never change when introducing an additional mutation at site *j*. This property allows us to define prior distributions where genetic interactions of order higher than |*U*| are allowed, but are constrained to specific sets of sites. Similarly, if we set all the 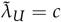 for *U < P* we see that the first sum in the exponent of Equation S53 becomes proportional to the squared norm of the projection of *f* into the null space of Δ^(*P*)^. Taking the limit *c* → ∞ is then equivalent to imposing a uniform prior on these directions, and for the special case *P* = 2 we recover Equation 4 in the main text.

Next, we consider the expected average squared epistatic coefficients involving sites *U* under the prior distribution *f* ~ N (0, *K*), parameterized by the lower-order variances 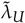 and *a*_*U*_ for |*U*| ≥ *P*. To draw samples from this prior, we can first sample *z* ~ N (0, *I*) and then set *f* = *K*^1/2^ *z*, so that Cov[*f*] = *K*^1/2^(*K*^1/2^)^T^ = *K* (for any matrix square root *K*^1/2^). One way to define the matrix square root is via its known eigendecomposition 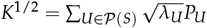. Then, we consider the average squared epistatic coefficient for any *f* and compute its expectation when *f* are drawn from the prior distribution as follows

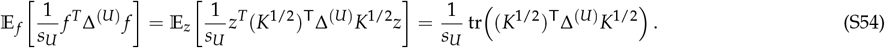

We compute the *K*^1/2^Δ^(*U*)^ *K*^1/2^ product by using the known eigendecompositions of *K* and Δ^(*U*)^:

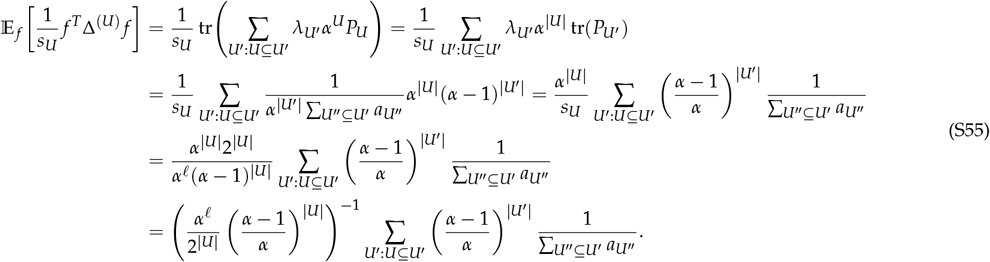

Thus, we can see that the expected average squared local epistatic coefficient for sites *U* does not depend only on the value *a*_*U*_, but also on all other *a*_*U*_′ such that both sets have at least one site in common (|*U* ∩ *U*′| *>* 0). For instance, if we let *a*_*U*_ → ∞ for a particular *U, λ*_*U*_′ for all *U*′ that include the whole set of sites in *U* will be set to zero, decreasing the expected average squared epistatic coefficients for sites involving sites in *U*′.

### Relationship with the Connectedness Model

In this section, we review the Connectedness Model and its relationship to Local Epistasis Regression. The Connectedness Model was first introduced as a Gaussian random field model by Reddy and Desai (2021) to allow different loci to have different probability of being involved in epistatic interactions with mutations at other sites. Then, it was used as a prior distribution in a Gaussian process model, uncovering the set of sites that are more strongly involved in epistatic interactions and using that information for inference of high-dimensional empirical fitness landscapes (Zhou et al. 2025). Here, we show that the Connectedness Model can be derived as a particular case of Eq. S48 as a function of the variance associated to the constant component 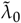 and the variance associated to the additive contribution of each site 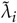

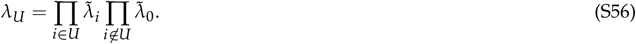

If we plug this into Eq. S48, we can see that the resulting kernel can be expressed as a product of site-specific kernels

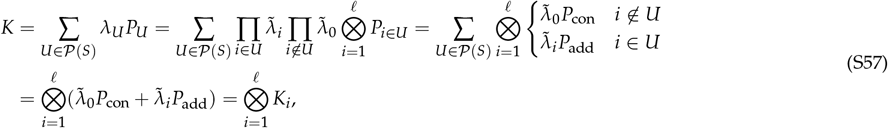

with entry-wise formula given by

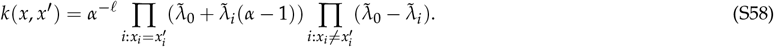

This kernel can be reparametrized as a function of the prior variance *σ*^2^ and the correlation under this kernel parametrized by 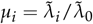 assuming 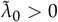 as in Zhou *et al*. (2025):

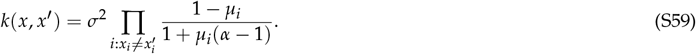

This construction shows that the variance explained by epistatic interactions under the Connectedness Model is completely specified by the variance explained by the additive contribution of each individual site. In contrast, Local Epistasis Regression allows arbitrary relationships between the additive and pairwise contribution of individual sites and pairs of sites, but the variance for higher-order interactions is fully specified by the the variance associated to pairwise interactions between specific pairs of sites (Eq. S52). These models also make different assumptions on how the variances associated to lower-order interactions combine to specify the variances associated to higher-order interactions. In particular, in the Connectedness Model these variances combine multiplicatively (Eq. S56), whereas in Local Epistasis Regression they are proportional to the harmonic mean of the variances associated to pairwise interactions between each possible pair of sites in *U*

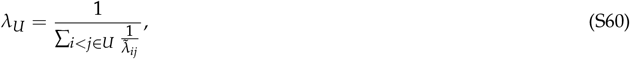

where 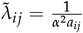

### Inference of fitness landscapes under the prior

In this section, we review how to do Gaussian process inference of a complete fitness landscape *f* from high throughput experimental data (Martí-Gómez et al. 2026b). We start by defining a Gaussian prior distribution over the space of possible fitness landscapes p(*f*) ~ *N* (0, *K*) that assigns higher probability to fitness landscapes that we believe are more plausible *a priori*.

Let *y* be an *n*-dimensional vector of measurements with known experimental Gaussian error given by the variance *n*-dimensional vector *y*_*var*_ for a subset of *n* ≤ *αℓ* sequences *X*. As both the prior distribution and the likelihood function are Gaussian, the posterior distribution is also multivariate Gaussian with closed form analytical solution (Rasmussen and Williams 2008) for the mean

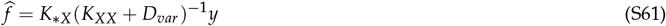

and covariance matrix

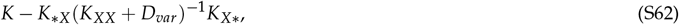

where *K*_*XX*_, *K* _* *X*_, *K* _* *X*_ are submatrices of *K* indexed by sequences *X* and *, where * represents all possible sequences, and *D*_*var*_ is a diagonal matrix with the known experimental variances *y*_*var*_ along the diagonal.

Despite the simplicity of the solution, practical evaluation of these expressions becomes challenging as the number of observations increases. Traditional approaches rely to computation of the Cholesky decomposition of the *K*_*XX*_ + *D*_*var*_ matrix to then use efficient triangular solves to compute the solutions to the linear systems rather than using direct matrix inversion for higher numerical stability. However, the algorithm for computing this decomposition is *O*(*n*^3^) and cannot be parallelized, which has traditionally limited the applicability of Gaussian process models to datasets with at most few thousand data points (Rasmussen and Williams 2008). In previous work, we have circumvented this limitation by leveraging the mathematical properties of the specific precision or kernel matrices, which could be expressed as polynomials in the Laplacian of the Hamming graph representing sequence space (Zhou and McCandlish 2020; Zhou et al. 2022; Chen et al. 2021; Martí-Gómez et al. 2026b). This property allowed us to encode these matrices as linear operators that allow computing matrix-vector products efficiently without explicitly storing them in memory, and use these operators together with iterative methods for solving systems of linear equations to scale these methods up to a few million data points.

Here we use a similar strategy by finding a representation of the kernel matrices that enables efficient computation of matrix-vector products without explicitly constructing them in memory, even if the kernel matrices presented here cannot be represented as polynomials in the Laplacian of the Hamming graph.

In the case of the Connectedness Model (Zhou et al. 2025), the kernel can in fact be expressed as a Kronecker product of *ℓ* site-specific *α* × *α* matrices (Eq. S57), which enables fast computation of matrix-vector products by leveraging the mixed-product property, which reduces the complexity to that of computing *ℓ* products of an *α* × *α* matrix with an *α* × *α*^*ℓ*−1^ matrix (Martí-Gómez et al. 2026b,a).

In the case of Local Epistasis Regression, the kernel matrix can be expressed as the sum of 2^*ℓ*^ matrices, each of which is Kronecker factorizable

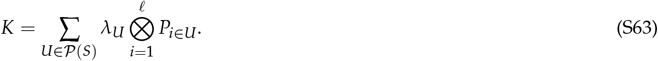

While this enables computation of matrix-vector products with a total of 2^*ℓ*^*ℓ α* × *α* by *α* × *α*^*ℓ*−1^ matrix products, the computational burden of these calculations is much larger compared with previous approaches e.g. requires 2^*ℓ*^ times the computation needed in the Connecntedness model. However, here we note that all of the 2^*ℓ*^ kernel matrices represent different combinations of only two different Kronecker factors *P*_con_ and *P*_add_, which allow us to re-use parts of the computation. In particular, we can decompose any *P*_*U*_ as follows

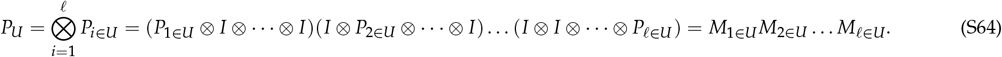

While the *M* matrices are still Kronecker products of *ℓ* matrices, *ℓ* − 1 of the factors correspond to the identity matrix and thus leave the matrices they act on unchanged, reducing the computation to a single *α* × *α* by *α* × *α*^*ℓ*−1^ matrix products. Importantly, *P*_*U*_ matrices differing only at the factor at the first site can be computed with a single additional operation from the same intermediate result

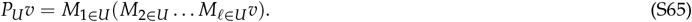

The intermediate results can also be computed in the same fashion

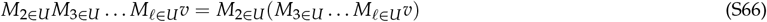

so that the computation can be shared with products of the same vector with other *P*_*U*_′ for a different subset of sites *U*′. These computational dependencies can be represented by a bifurcating tree where nodes represent *α*^*ℓ*^-dimensional vectors and edges represent matrix-vector products with specific *M*_*i* ∈ *U*_ matrices. The vector *v* is located at the root of the tree, allowing computation of the intermediate vectors at each of the nodes of the tree up to the tips containing all the *P*_*U*_*v* for every possible *U*. This algorithm reduces the total number of operations from 2^*ℓ*^*ℓ* to 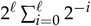. This series converges relatively fast to 2^*ℓ*+1^, resulting in an approximate *ℓ*/2-fold speedup even when *ℓ* is small. For instance, for *ℓ* = 8, the number of matrix-matrix products goes from 8 × 2^8^ = 2048 under the naive approach to 510, nearly achieving the expected 4-fold increase in computational efficiency.

### Hyperparameter optimization

In order to infer a fitness landscape under a given prior distribution, we must first choose the parameters that define the properties of the prior, also known as hyperparameters. Here, we generally consider prior distributions defined over the space of possible fitness landscapes *f* defined by a kernel function *k*(*x, x*′) that returns the covariance for any pair of sequences *x* and *x*′ given by

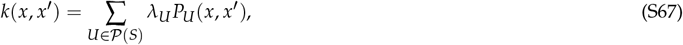

where *P*_*U*_ (*x, x*′) is the covariance due to interactions between exactly *U* sites between sequences *x* and *x*′, which depends only on the combination of sites at which they differ *P*_*U*_ (*x, x*′) = *w*_*U*_ (*H*(*x, x*′)); and where the parameters *λ*_*U*_ can be free or a function of a smaller set of parameters generally called *θ* (*λ*_*U*_ = *g*(*θ*)) e.g. Eq. S52 in Local Epistasis Regression and Eq. S56 in the Connectedness Model.

In this work, we use a strategy known as kernel alignment (Wang et al. 2015; Zhou et al. 2022) or Haseman-Elston regression (Haseman and Elston 1972), in which the prior covariance approximates as closely as possible the patterns of covariance in the empirical data. Specifically, this is done by finding the parameter values 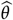 that minimize the Frobenius norm of the difference between the prior predictive covariance *K*_*XX*_ + *D*_*var*_ and the empirical second moment matrix *yy*^*T*^ :

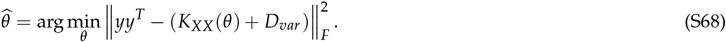

Naively solving this minimization problem is challenging, as we need to work with *n* × *n* matrices, where the number of measured sequences *n* can be in the order of hundreds of thousands to millions. However, as the prior covariance between two sequences only depends on the set of sites at which they differ, the dimensionality of the problem can be reduced to a more manageable 2^*ℓ*^-dimensional weighted least squares problem as follows:

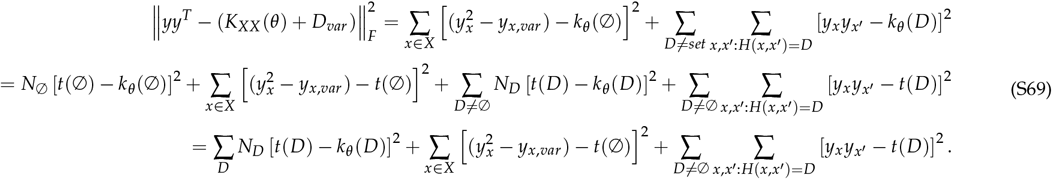

Since the last two terms are independent of *θ*, we can find by optimal hyperparameter values 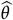 simply as

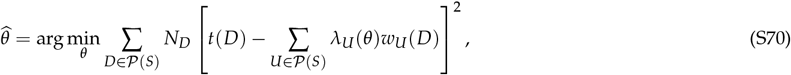

where *t*(*D*) corresponds to the second moment, related with the empirical autocovariance function 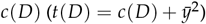 given by

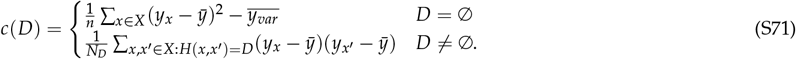

## Supplementary figures

**Figure S1.**
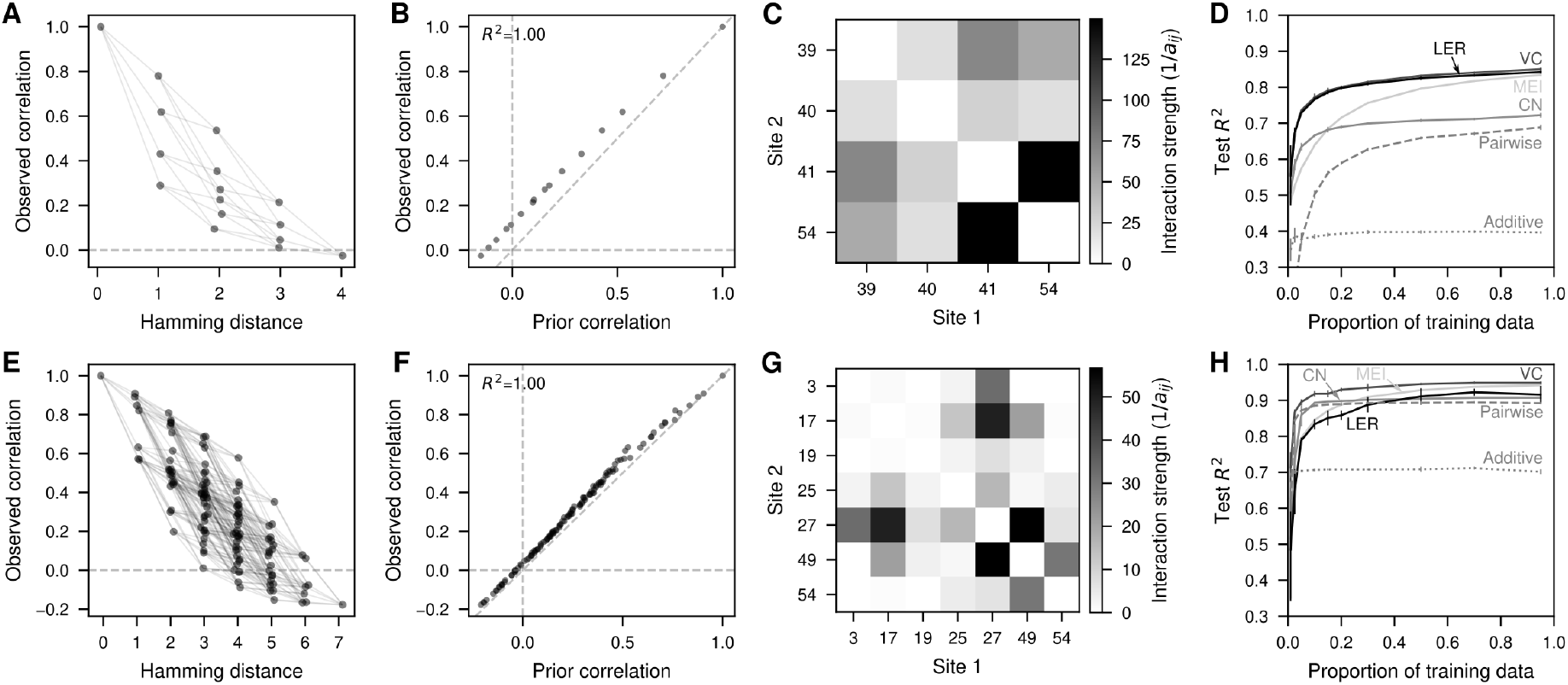
Application of Local Epistasis Regression to protein datasets. (A,E) Correlation in the measured fitness values for pairs of sequences differing at each possible subset of sites *D* arranged according to the Hamming distance *d* = |*D*|. Each dot represents a single distance class *D* and are joined by lines whenever the distance classes differ by a single position from each other. (B,F) Comparison of the observed correlation values in the data and the values under the estimated prior ones using Local Epistasis Regression for every possible distance class *D* (each dot represents a different *D*). Correlations were estimated using 80% of the data for training. (C,G) Heatmap representing the inferred model hyperparameters as 1/*a*_*ij*_ for every pair of sites *i, j* highlighting the patterns of genetic interactions across sites under the prior. (D,H) Predictive performance evaluated by the *R*^2^ between the predicted and the measured fitness of held-out test sequences when using different amounts of training data for different models (MEI: Minimum Epistasis Interpolation, VC: Variance Component regression, CN: Connectedness Model regression, LER: Local Epistasis Regression). Predicted values are the maximum a posteriori estimate given by each method, which is equal to the posterior mean *f*. Error bars represent the standard deviation across 3 different random samples for each fraction of training data. Each row represents a fitness landscape: GB1 (A,B,C,D); FYN-SH3 (E,F,G,H).

**Figure S2.**
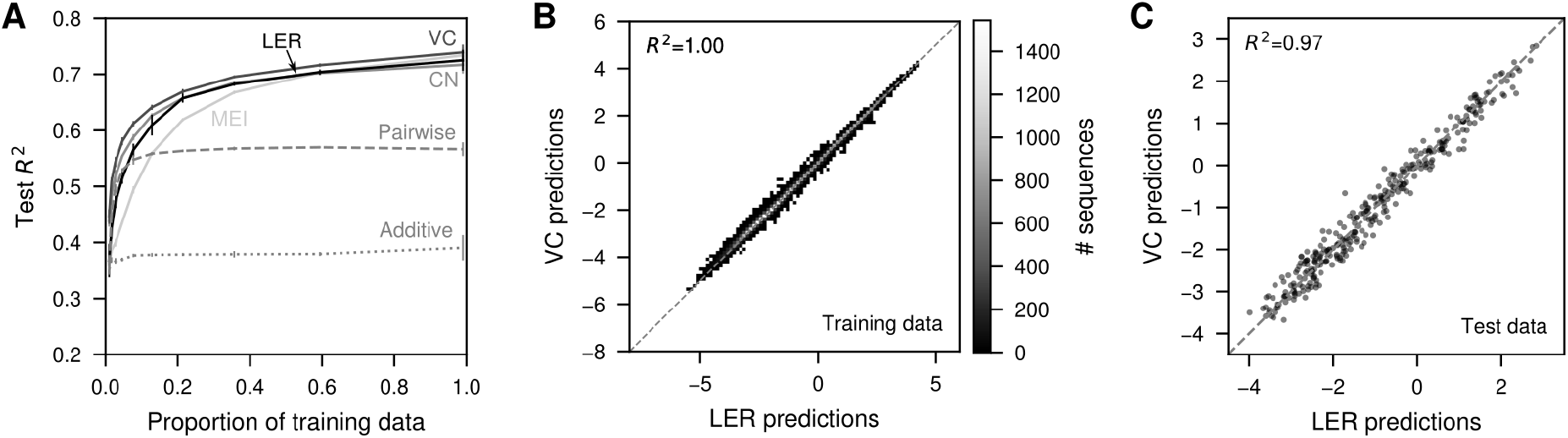
Comparison of model predictions on the self-splicing intron dataset. (A) Predictive performance evaluated by the *R*^2^ between the predicted and the true fitness values of held-out test sequences when using different amounts of training data for different models (MEI: Minimum Epistasis Interpolation, VC: Variance Component regression, CN: Connectedness Model regression, LER: Local Epistasis Regression). Predicted values are the maximum a posteriori estimate given by each method, which is equal to the posterior mean *f*. Error bars represent the standard deviation across 3 different random samples for each fraction of training data. (B) 2D histogram comparing the fitness landscape reconstructions of the self-splicing intron dataset under Local Epistasis Regression (LER) and Variance Component regression (VC). (C) Scatterplot comparing the predictions in held-out sequences of the self-splicing intron dataset under Local Epistasis Regression (LER) and Variance Component Regression (VC).

**Figure S3.**
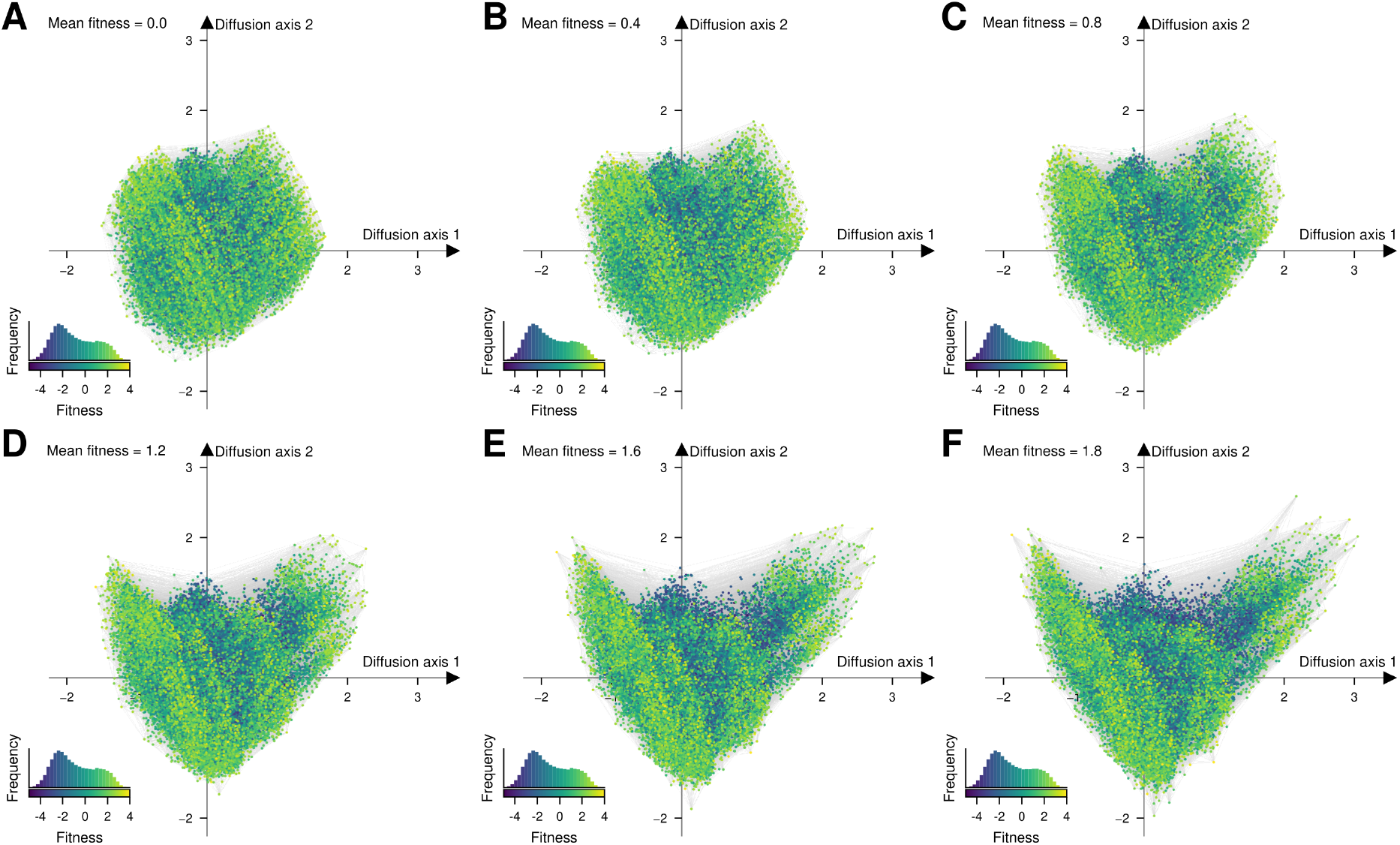
Visualization of the inferred fitness landscape using Local Epistasis Regression under different strengths of selection, where here we quantify the strength of selection by the mean fitness achieved at stationarity, i.e. under long-term purifying selection. Every dot represents one of the possible 4^8^ possible sequences and is colored according to the predicted fitness. The inset represents the phenotypic distribution along with their corresponding color in the map. Sequences are laid out according to the first two Diffusion axes and dots are plotted in order according to Diffusion axis 3.

**Figure S4.**
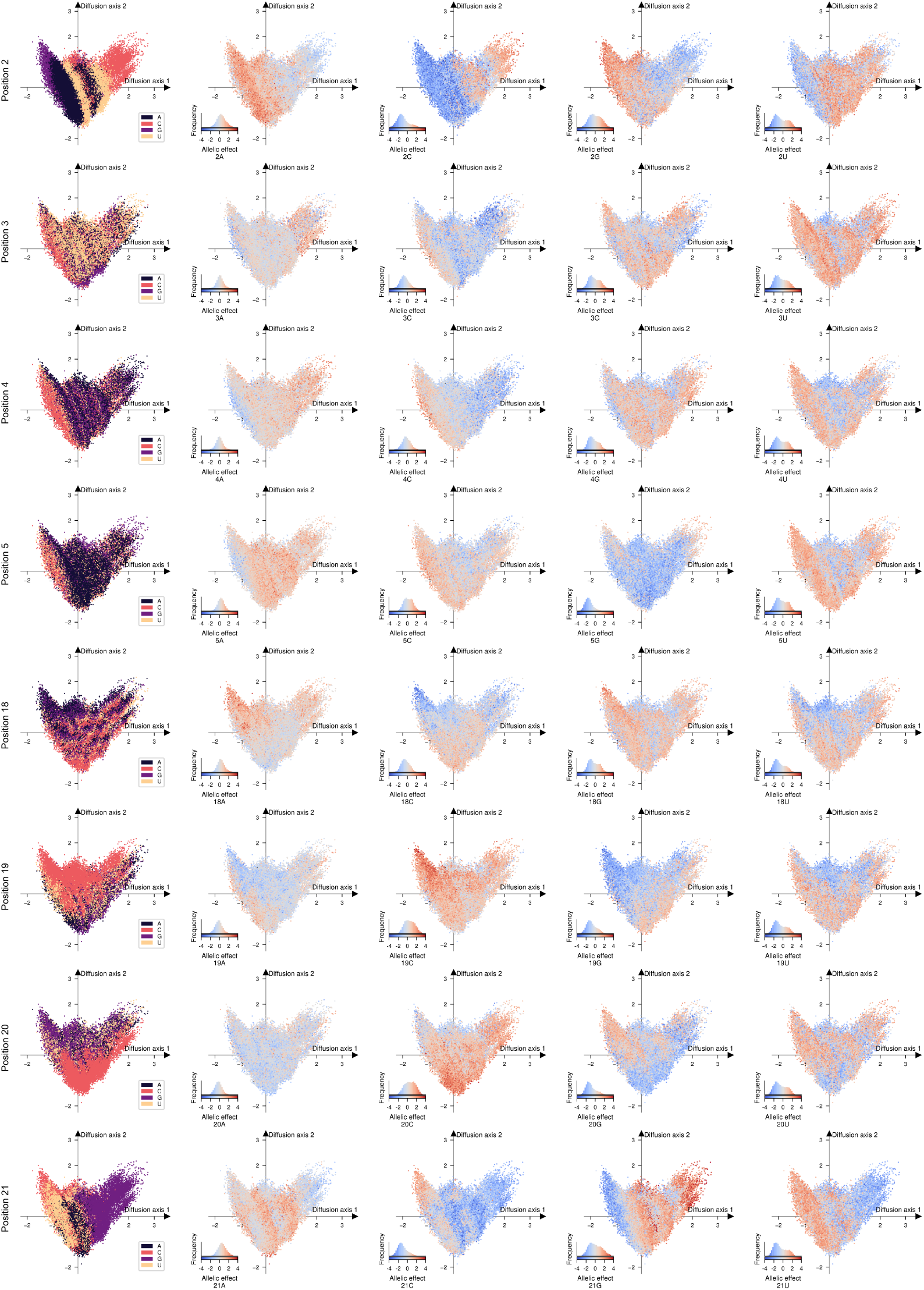
Visualizing alleles and allelic preferences across the fitness landscape visualization. Visualization of the inferred fitness landscape using Local Epistasis Regression. Every dot represents one of the possible 4^8^ possible sequences and is colored according to the allele (first column) or the difference in the fitness of the sequence obtained when placing a specific allele at an specifc position relative to the average fitness of the four possible alleles (four last columns). The inset represents the allelic effect distribution along with their corresponding color in the map. Sequences are laid out according to the first two Diffusion axes and dots are plotted in order according to Diffusion axis 3.

## Notes

https://github.com/cmarti/deltaU

### Competing Interest Statement

The authors have declared no competing interest.

